# Neural tube organoids self-organise floorplate through BMP-mediated cluster competition

**DOI:** 10.1101/2023.06.25.546258

**Authors:** Teresa Krammer, Hannah T. Stuart, Elena Gromberg, Keisuke Ishihara, Manuela Melchionda, Jingkui Wang, Elena Costantini, Stefanie Lehr, Dillon Cislo, Laura Arbanas, Alexandra Hörmann, Ralph A. Neumüller, Nicola Elvassore, Eric Siggia, James Briscoe, Anna Kicheva, Elly M. Tanaka

## Abstract

The neural tube (NT) has been a hallmark example of embryonic induction and patterning whereby the notochord induces an organiser, the floorplate, that secretes Sonic Hedgehog (SHH) to pattern the surrounding field of neural progenitors. On the other hand, NT organoids (NTOs) formed from embryonic stem cells (ESCs) undergo spontaneous floorplate formation and patterning in the absence of their normal embryonic inducers. Understanding how stem cells undergo regulative organiser formation is a central challenge in biology. Here, we investigated the self-organisation of a SHH-expressing floorplate organiser using clonal NTOs. Expression of FOXA2, a floorplate transcription factor, was initially spatially scattered before resolving into multiple clusters. These FOXA2^+^ clusters underwent competition and physical sorting, resulting in a stable “winning” floorplate. We identified BMP signalling as a key governor of long-range cluster competition. FOXA2^+^ clusters expressed BMP4 ligand suppressing FOXA2 in receiving cells, while simultaneously expressing the BMP-inhibitor NOGGIN to secure FOXA2^+^ cluster survival. Genetic mutation of *Noggin* perturbed the floorplate not only in NTOs but also *in vivo* at the mid-hindbrain region of the mouse NT. These results demonstrate how the floorplate can form autonomously without its well-known inducer, the notochord, suggesting redundant mechanisms ensuring robustness. Defining molecular pathways that govern organiser self-organisation is critical in harnessing the developmental plasticity of stem cells toward directed tissue engineering.

## Introduction

During vertebrate development groups of cells called organisers secrete morphogens to induce cell fate patterning in the surrounding tissues ^1, 2^. In the embryo, organisers are induced by interaction of emerging tissues (reviewed in ^3^). Correct organiser placement relies on pre-existing asymmetries in the embryo and the delivery of polarising cues. Intriguingly, recent discoveries *in vitro* have found that organisers can form by a process of self-organisation, in the absence of their natural developmental inducers ^4–7^. Despite their central importance for multicellular development, as well as tissue regeneration and engineering, the mechanisms by which organisers self-organise are poorly understood.

The neural tube (NT) is the embryonic precursor of the central nervous system. Patterning of neural progenitor identities along the body axes ensures the formation of the correct diversity and arrangement of neurons ^8, 9^. In the developing spinal cord, NT patterning along the dorsal-ventral axis is driven by two opposing organisers: the dorsal roofplate secreting BMPs and WNTs, and the ventral floorplate secreting SHH morphogen ^10–13^. *In vivo*, the ventral floorplate organiser is usually induced by SHH and other signals from the underlying notochord ^14, 15^, which has been most extensively studied at thoracic levels. The more anterior floorplate is induced earlier during amniote gastrulation ^14, 16, 17^ and forms independently of notochord signalling by a mechanism that is currently unclear.

Remarkably, SHH-expressing floorplate has been shown to self-organise in NT organoids (NTOs) *in vitro* ^6^ in the absence of notochord or any directional inductive cues. NTOs form clonally, by differentiating single mouse embryonic stem cells (ESCs) embedded in 3D hydrogels ^6, 18^, resulting in neuroepithelial cysts with a single apical lumen that exhibit a “default” dorsal midbrain identity. A global pulse of retinoic acid (RA) applied at Day 2 for 18 hours posteriorises NTOs to hindbrain levels and triggers self-organisation of dorsal-ventral patterning by Day 6 ^6^ (Figure 1A). This phenomenon is characterised by the formation of a localised FOXA2^+^SHH^+^ floorplate at the emergent ventral pole, which then drives ventral-to-dorsal patterning of neural progenitor identities. Here, we address the mechanism governing the self-organisation of this floorplate organiser in NTOs.

**Figure 1:**
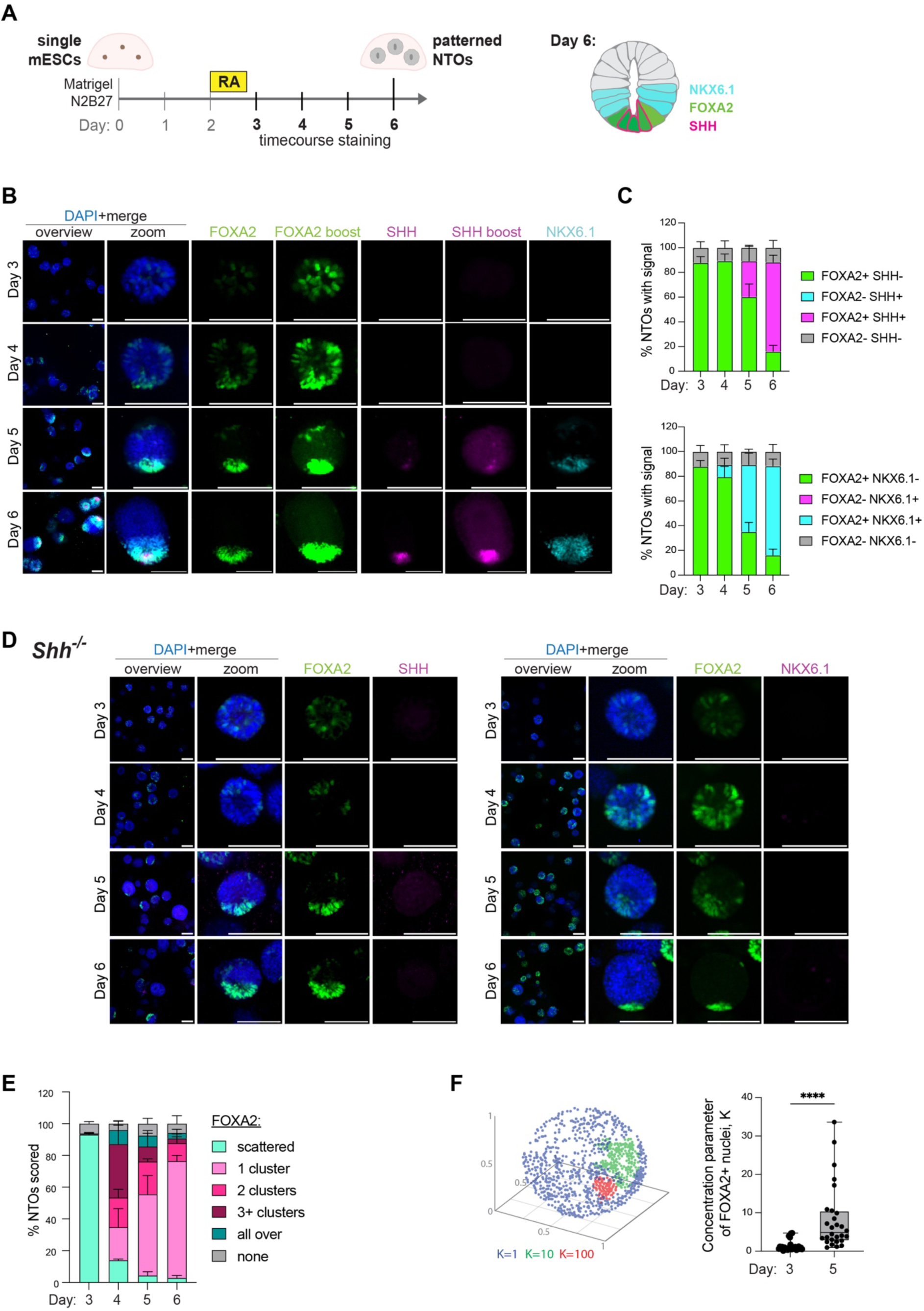
Clonal NTOs transition from a scattered expression of FOXA2+ cells to a clustered state which can occur in the absence of SHH. **A:** Schematic of NTOs formation. Single embryonic stem cells form epithelial NTOs within two days in Matrigel and N2B27. An 18-hour pulse of retinoic acid induces floorplate formation and DV patterning at the endpoint Day 6. **B:** IF timecourse of NTOs fixed from Day 3-6 showing (from left to right, single *z* slice) a merged overview image at 10x magnification, and a representative NTO (zoom) stained for DAPI (blue) and the DV patterning markers FOXA2 (green), SHH (magenta), NKX6.1 (cyan). In the second FOXA2 column, FOXA2 images were scaled with lower maximal grayscale value to highlight the early spatial pattern. In the second SHH column, the same metric was applied. Scale bars, 100 µm. **C:** Quantification of marker expression in (*n* = 528) NTOs over time. Coloured bars correspond to staining in B (*n* = 192 for Day 3, *n* = 158 for Day 4, *n* = 134 for Day 5 and *n* = 44 for Day 6). **D:** *Shh^-/-^* NTOs undergo FOXA2-cell clustering. Timecourse of *Shh^-/-^* NTOs fixed from Day 3-6 showing (from left to right, single *z* slice) a merged overview image at 10x magnification, and a representative NTO (zoom) in stained for DAPI (blue) and the DV patterning markers FOXA2 (green) and SHH or NKX6.1 (magenta). For parental control, see Figure S1. Scale bars, 100 µm. **E:** FOXA2^+^ cell cluster distribution over time. FOXA2 spatial distribution was classified into “scattered”, “1 cluster”, “2 clusters”, “3 or more clusters”, “none” (FOXA2^-^), or “all over” (almost entirely FOXA2^+^). A total of 1148 NTOs were manually analysed (*n* = 292 for Day 3, *n* = 285 for Day 4, *n* = 209 for Day 5 and *n* = 362 for Day 6). **F:** FOXA2^+^ cell clustering analysis using von Mises–Fisher parameter. Left: Schematic illustrating quantitative clustering parameter (K, from von Mises–Fisher). Right: von Mises– Fisher analysis of 3D FOXA2^+^ cell distribution in Day 3 (*n* = 27) and Day 5 (*n* = 28) NTOs. By Day 5, FOXA2-expressing cells show a higher K value. Each dot represents the K value of a single NTO. Results were statistically analysed using Mann-Whitney test with *p* = <0.0001. Error bars show standard deviation (SD), asterisks mark statistical significance.

## Results

### FOXA2^+^ floorplate precursors self-organise from a spatially scattered to clustered state in clonal NTOs

To characterise the spatiotemporal dynamics of how the FOXA2^+^SHH^+^ floorplate self-organises in NTOs, we first performed timecourse immunofluorescence (IF) staining after RA pulse (Figure 1A). We assayed FOXA2, SHH and SHH/GLI-responsive NKX6.1 (Figure 1B) and quantified the number of NTOs expressing each marker (Figure 1C). Temporally, FOXA2 expression was observed from Day 3 onwards, in >80% of NTOs, prior to detection of SHH or NKX6.1 proteins. From Day 5-6, the number of NTOs with FOXA2 and detectable SHH/NKX6.1 increased, whereas SHH or NKX6.1 expression were not observed in FOXA2^-^ NTOs (Figure 1C), indicating that FOXA2 preceded SHH expression and activity. Accordingly, RNA-seq analysis ^19^ showed upregulation of *FoxA2* transcript earlier than *Shh* or *Nkx6.1* (Figure S1A). IF and RNA-seq both showed an increase in the level of FOXA2 expression over time from Day 3 onwards (Figure 1B, S1A).

*In vivo*, SHH from the notochord can induce floorplate FOXA2 expression via GLI binding sites found in FOXA2 regulatory elements ^15, 20^. Although we observed FOXA2 expression prior to SHH in NTOs (Figure 1B-C, S1A), we nonetheless asked whether genetic deletion of SHH would abolish FOXA2 induction and cluster formation. NTOs formed from *Shh^-/-^*ESCs showed no detectable SHH or NXK6.1 expression at any timepoint (Figure 1D, S1B). However, FOXA2 induction and cluster formation still occurred in *Shh^-/-^* NTOs (Figure 1D, S1C). Unlike parental wildtype (WT) NTOs, the proportion of *Shh^-/-^*NTOs exhibiting FOXA2 expression decreased from 80% to 60% over Days 5–6 (Figure S1B), suggesting a role for SHH in FOXA2 maintenance at later timepoints, consistent with phenotypes of prolonged SHH antagonism in NTOs ^6^ and the SHH-driven regulatory architecture at the *FoxA2* locus ^20^.

Spatially, NTOs expressed FOXA2 at Day 3 in a scattered salt-and-pepper manner (Figure 1B). Subsequently, the FOXA2 signal became increasingly clustered, until by Day 5-6 FOXA2 was found in discrete domains, inside which SHH expression commenced to yield a functional floorplate reflected by induction of NKX6.1 (Figure 1B-C). Over time, the proportion of NTOs with scattered FOXA2, versus 3+, 2 or 1 clusters per NTO shifted until a majority of NTOs contained a single FOXA2^+^ cluster (Figure 1E). To quantify the spatial distribution of FOXA2^+^ nuclei in 3D, we performed von Mises-Fisher analysis in which a concentration parameter *K* is computed based on the sum of directional vectors from the NTO centre. This confirmed that FOXA2 expression became increasingly concentrated over time (Figure 1F). Together, this shows that floorplate self-organisation in RA-treated NTOs is characterised by “salt-and-pepper” FOXA2 induction, followed by clustering of FOXA2^+^ floorplate precursor cells, which occurs prior to and independently of SHH expression.

### Intermediate FOXA2^+^ clusters interact via competition and sorting during self-organisation

We next sought to understand whether and how individual NTOs progress from scattered to multi-cluster to single floorplates. We generated a FOXA2-Venus fusion reporter, together with knock-in of H2A-mCherry into the safe-harbour *Rosa26* locus to constitutively label nuclei (Figure 2A, S2A-C). Using 3D light-sheet live-imaging, we acquired images every 15 minutes starting from Day 3 in over 300 live NTOs (Figure 2B). This revealed self-organisation of FOXA2-Venus^+^ cells from a spatially scattered to a clustered state via progressive formation of and resolution between intermediate clusters (Figure 2C, Movies S1-2), in agreement with fixed sample quantifications (Figure 1E).

**Figure 2:**
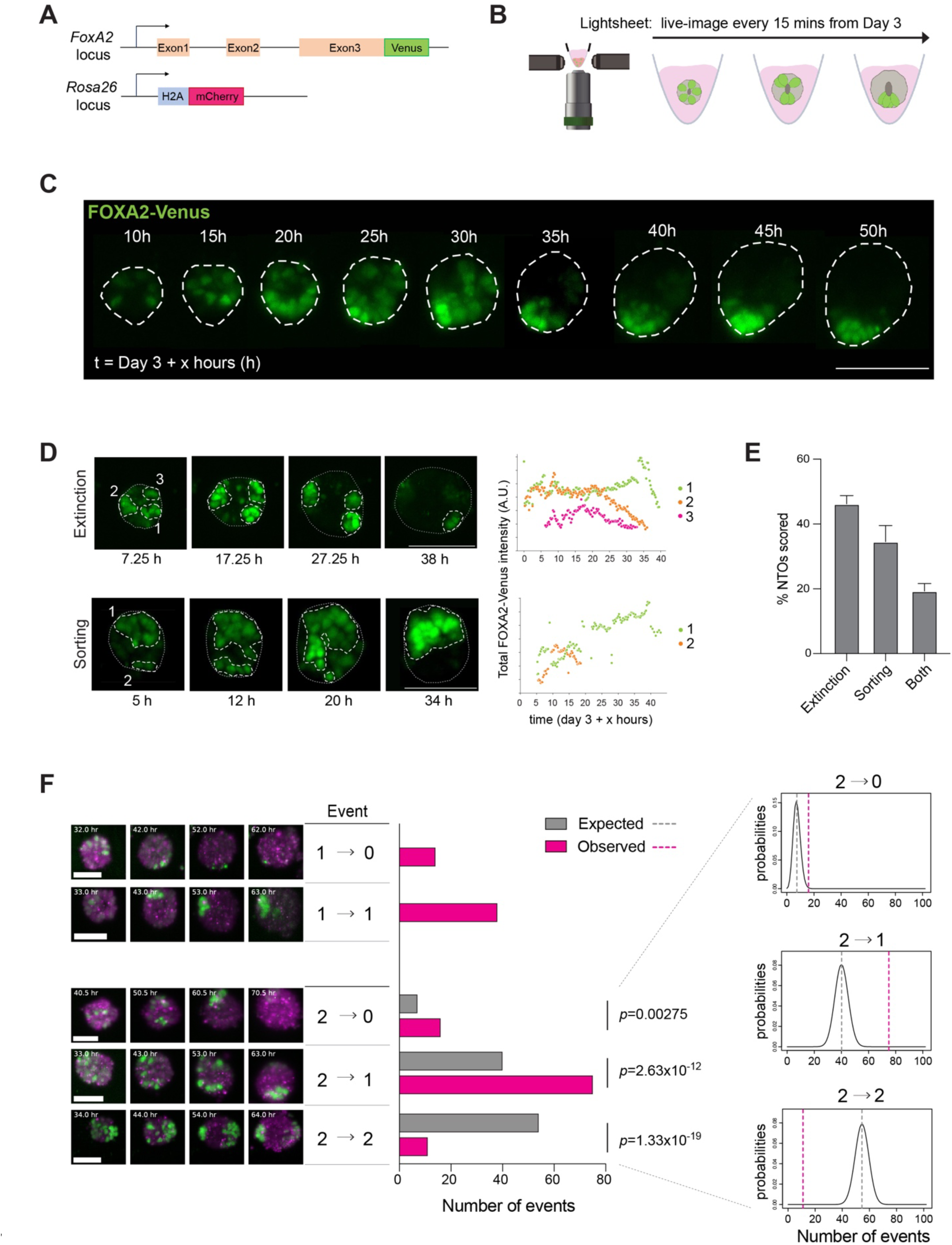
Statistical analysis of live cluster dynamics reveals cluster competition during self-organisation. **A:** Schematic of double reporter generation. Venus was knocked in after the 3^rd^ exon at the C-terminus of the endogenous FOXA2 locus generating a fusion protein. In addition, an H2A-mCherry construct was knocked into the ubiquitously expressed Rosa26 locus to mark nuclei. **B:** Schematic (created with BioRender) of timelapse imaging of double reporter on the Viventis lightsheet microscope. Full image stacks with 3 µm *z* step size were taken every 15 mins starting 6 hours post RA removal on Day 3 for 48-75 hours. **C:** Images from timelapse series of a NTO (maximum intensity *z*-projections) following the spatial organisation of FOXA2 (green) over 50 hours showing self-organisation from a dispersed (salt-and-pepper) into a clustered state (see Movies S1-2). Dashed white lines mark the outlines of the NTO. Scale bar, 100 µm. **D:** Two distinct modes of cluster formation. Selected maximum intensity *z*-projection images from two (upper and lower) NTO timelapse series. Upper: “Extinction”: three FOXA2^+^ (green) clusters form, and then one remains while two disappear (see Movies S3-4). Lower: “Sorting”: Two distinct clusters grow and coalesce into a single cluster over time (see Movies S5-6). Right: Cluster intensities were tracked over time in graphs. Dotted grey lines mark the outlines of NTOs, dashed white lines mark the outline of FOXA2^+^ clusters. Scale bars, 100 µm. **E:** A total of *n* = 93 NTOs from three different timelapses that displayed as 2è1 behaviour were subclassified into their mode of self-organisation (extinction, sorting, or both), taking into account that sorting might be overestimated due to analysis using maximum intensity *z*-projections. **F:** Left: representative timecourse images for each category shown as maximum intensity *z-* projections, scale bars 50 µm, and scoring of NTOs according to FOXA2-Venus^+^ cluster numbers at intermediate vs endpoint times. Middle: based on the frequency of cluster disappearance in NTOs with one intermediate cluster (*n* = 52), the expected outcomes (grey) for NTOs with 2 intermediate clusters (*n* = 102) were calculated using binomial distribution assuming independent cluster behaviour, then compared to observed outcomes (magenta). Right: calculation of *p* values based on the binomial distribution for each case, with the expected value indicated by grey lines and the observed numbers by magenta lines.

We observed two distinct modes by which intermediate clusters resolved (Figure 2D, Movies S3-6): 1) cluster extinction, whereby FOXA2-Venus expression was lost in some clusters whilst it persisted in others, and 2) physical cell-sorting, whereby FOXA2-Venus^+^ cells from one cluster coalesced with another cluster. These two behaviours were not mutually exclusive: 20% of NTOs exhibited both sorting and extinction of FOXA2-Venus^+^ clusters, whilst 35% and 45% exhibited only sorting or only extinction, respectively (Figure 2E). We focussed on cluster extinction as it was observed in the majority of NTOs.

To understand if cluster extinction occurred autonomously or via communication (competition) between clusters, we categorised live-imaged NTOs according to the number of FOXA2-Venus^+^ clusters observed at intermediate versus endpoint times. We then tested statistically whether clusters behaved independently of each other. For NTOs with one intermediate cluster, 14 out of 52 extinguished their cluster during self-organisation (Figure 2F). By considering the disappearance of a given cluster as a process with this observed probability, we formulated the null hypothesis that, if FOXA2-Venus^+^ clusters behave independently, the outcome for NTOs with two intermediate clusters can be explained by multiplying the probabilities. We thus calculated the expected outcomes then compared them with experimental observations (Figure 2F). In live-imaged NTOs with two intermediate clusters, the observed outcomes differed significantly from the expected outcomes. A cluster was significantly more likely to be extinguished if another cluster was present (Figure 2F), indicating competitive communication between the clusters that governs final outcome.

### BMP signalling activity anti-correlates with FOXA2 expression during cluster interaction

We sought to determine the molecular signals that facilitate cluster communication. To identify candidates, we performed bulk RNA-seq of sorted FOXA2-Venus positive versus negative cells at Day 5, a timepoint during which cluster extinction was occurring (Figure 3A, S3A). Reasoning that secreted signals could explain long-range mutual inhibition between clusters, we focused on the expression of ligands, receptors, transducers and modulators for the major secreted signalling pathways. Using two methods to test enrichment of signalling pathway signatures in FOXA2^+^ versus FOXA2^-^ cells (Figure 3B, S3B), signatures of WNT, FGF and BMP signalling were uncovered in FOXA2^+^ cells. Various WNT and FGF ligands were expressed in FOXA2^+^ or FOXA2^-^ populations (Figure 3C). Notably, BMP ligands were almost exclusively expressed in FOXA2^+^ cells (Figure 3C). Expression of the secreted WNT inhibitors *Dkk1-3* and secreted BMP inhibitors *Noggin* and *Chordin* was enriched in FOXA2^+^ cells, whereas FGF-responsive *Spry4* was enriched in FOXA2^−^ cells (Figure 3C). We therefore investigated WNT, FGF and BMP pathways in NTOs, by spatial analyses of signalling activities and functional perturbations as described below.

**Figure 3:**
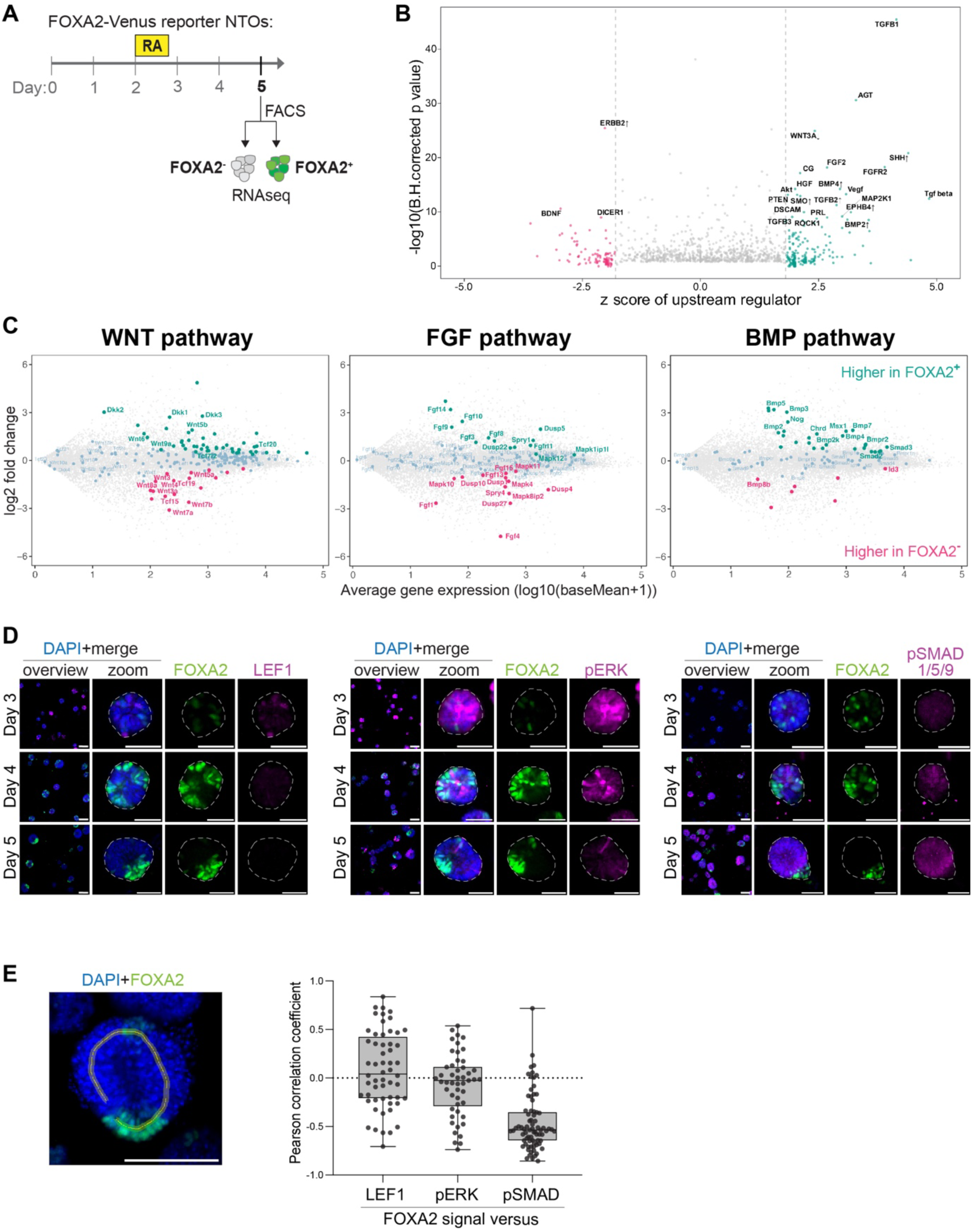
FOXA2+ cells express *BMP*s and *Noggin* while FOXA2-cells display higher pSMAD 1/5/9 signal. **A:** Schematic of RNA-seq performed on sorted FOXA2^+^ and FOXA2^-^ cell populations of Day 5 NTOs made from the reporter cell line. **B:** Activity plot of upstream regulators predicted to be activated or inhibited based on the observed expression changes in FOXA2^+^ versus FOXA2^-^ cells. Regulators are coloured based on activation Z-score at a threshold of 1.8, and the top 20 candidates based on significance are labelled; an arrow next to a regulator name indicates significant up or down regulation of the gene in FOXA2^+^ (vs FOXA2^-^). **C:** WNT, FGF and BMP pathway members showing differential gene expression between FOXA2^+^ and FOXA2^-^ cells. MA-plot of the RNA-seq data showing log2 fold changes of genes over their average gene expression against samples (*y* axis: FOXA2^+^ versus FOXA2^-^ log ratio values, *x* axis: log10(baseMean+1) of average gene expression). Colour highlights genes assigned to either WNT signalling pathway (GO:0016055), FGF signalling pathway (GO:0008543), or BMP signalling pathway (GO:0030509) based on Ensembl 94 annotation. Data point colour indicates up-regulated in FOXA2^+^ (green, #009988), up-regulated in FOXA2^-^ (pink, #EE3377) or not significantly differentially regulated genes (light blue). **D:** Time course of NTOs fixed from Day 3-5 (single *z* slice) showing a merged image at 10x magnification, and a representative NTO (zoom) stained for DAPI (blue), FOXA2 (green) and LEF1 to visualise WNT (left)/ pERK to visualise FGF (middle)/ pSMAD1/5/9 to visualise BMP pathway activity (right) (magenta). Dashed white lines mark the outlines of the NTOs. Scale bars, 100 µm for 10x, and 50 µm for zoom of representative NTOs. **E:** Analysis of Pearson correlation coefficient. Left: Representative NTO used for the anti-correlation analysis performed on single *z* slices of images on Day 5. For each NTO cross-section, a line with width of 10 pixels (shown in yellow) was drawn around the NTO starting in the FOXA2^+^ cluster sparing the lumen. Scale bar, 100 µm. Right: Intensity profiles of FOXA2 and LEF1/ pERK/ pSMAD were measured and analysed by calculating the Pearson correlation coefficient by sampling the pixels for two channels (FOXA2 – LEF1, FOXA2 – pERK, FOXA2 – pSMAD). NTOs are *n* = 56 LEF1, *n* = 49 pERK, *n* = 76 pSMAD.

To functionally test the candidate pathways, we performed pharmacological screens using small molecules and recombinant proteins to activate or inhibit WNT, FGF or BMP signalling from Day 3-5 (Figure S3C). WNT or FGF perturbations led to a slight decrease in the percentage of FOXA2^+^ NTOs at Day 5, irrespective of whether the pathway was activated or inhibited. However, BMP activation decreased while inhibition increased the percentage of FOXA2^+^ NTOs, demonstrating that the level of BMP signalling modulates FOXA2 expression.

To assess the activities of the WNT, FGF and BMP pathways during cluster self-organisation in NTOs, we assayed for LEF1, pERK or pSMAD1/5/9 respectively by IF (Figure 3D). LEF1 and pERK signal decreased whereas pSMAD1/5/9 increased from Day 3-5, suggesting an increase in BMP pathway activity after RA treatment. Interestingly, we observed that pSMAD1/5/9 signal predominated in FOXA2^-^ cells compared to FOXA2^+^ cells (Figure 3D, S3D-E). To measure the degree of spatial correlation between FOXA2 expression versus WNT, FGF or BMP signalling activities, we measured signal intensities along a line that bisected cells in NTOs, circling the lumen, then computed the Pearson correlation coefficients between FOXA2 and LEF1, pERK or pSMAD1/5/9 signals respectively (Figure 3E). We did not observe any type of correlation between FOXA2 and LEF1 or pERK signal. In contrast, a negative correlation was found between FOXA2 and pSMAD1/5/9 intensities, consistent with a role of BMP signalling in FOXA2 suppression. Taken together, the BMP pathway exhibited an appropriate activity pattern and perturbation phenotype to be further considered as a candidate signal by which intermediate FOXA2 clusters interact.

### BMP signalling regulates FOXA2 cluster resolution

To functionally investigate the role of BMP signalling during the phase of cluster competition, we perturbed the BMP pathway from Day 4 onwards using pharmacological and genetic approaches (Figure 4A). To quantitatively assess the effect of perturbations on inter-cluster communication, we adapted our protocol to support the growth of larger NTOs (Figure S4A-C) that correspondingly demonstrated a higher frequency of multiple rather than single FOXA2^+^ clusters (Figure S4D) due to a positive correlation between NTO size and endpoint cluster number (Figure S4E).

**Figure 4:**
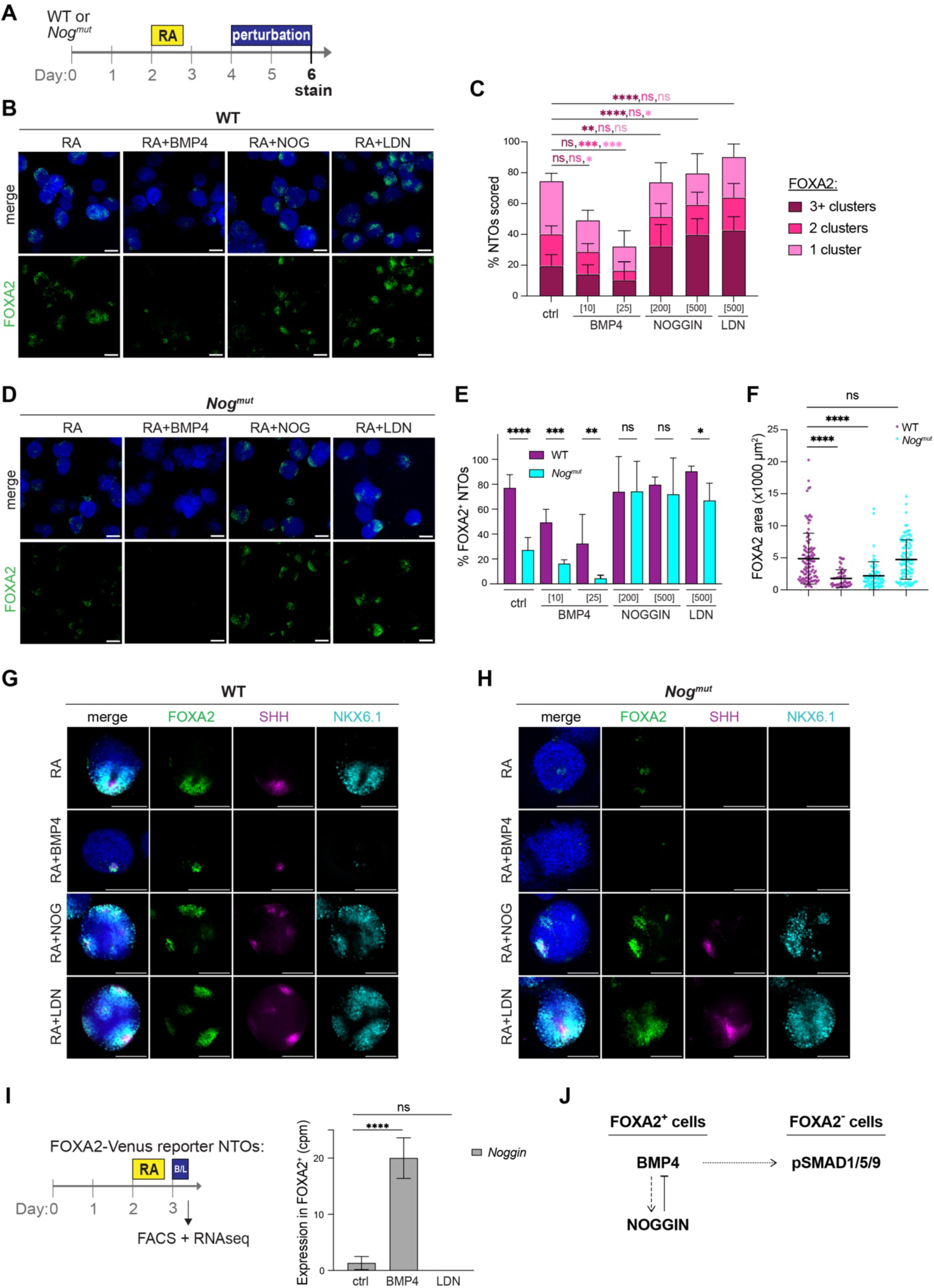
BMP pathway activation or suppression modulates FOXA2 cluster persistence. **A:** Timeline of BMP pathway perturbations using “large NTO” protocol relevant for panels B-H (see Figure S4). Perturbations were performed from Day 4-6 after an 18-hour RA pulse from Day 2-3. **B:** Representative images at 10x magnification (single *z* slice) of WT NTOs stained for DAPI (blue) and FOXA2 (green) for RA, RA+BMP4 (25 ng/ml), RA+NOG (200 ng/ml) and RA+LDN (500 nM). Scale bars, 100 µm. **C:** 3D analysis of numbers of FOXA2^+^ clusters in WT NTOs on Day 6 shown in B. Results were statistically analysed using an ordinary one-way ANOVA and Dunnettes’s multiple comparisons test. For 3 or more clusters: RA vs. RA+BMP4 (10 ng/ml) was n.s., RA vs. RA+BMP4 (25 ng/ml) was n.s, RA vs. RA+NOG (200 ng/ml) *p* = 0.0029, RA vs. RA+NOG (500 ng/ml) *p* = <0.0001, RA vs. RA+LDN (500 nM) *p* = <0.0001; for 2 clusters: RA vs. RA+BMP4 (10 ng/ml) was n.s., RA vs. RA+BMP4 (25 ng/ml) *p* = 0.0005, RA vs. RA+NOG (200 ng/ml) was n.s., RA vs. RA+NOG (500 ng/ml) was n.s., RA vs. RA+LDN (500 nM) was n.s.; for 1 cluster: RA vs. RA+BMP4 (10 ng/ml) *p* = 0.0171, RA vs. RA+BMP4 (25 ng/ml) *p* = 0.0007, RA vs. RA+NOG (200 ng/ml) was n.s., RA vs. RA+NOG (500 ng/ml) *p* = 0.0153, RA vs. RA+LDN (500 nM) was n.s. Concentration [x] expressed in ng/ml for BMP4 and NOGGIN, or in nM for LDN. Asterisks indicate statistical significance, n.s. indicates non-significance. **D:** Representative images at 10x magnification (single *z* slice) of *Nog^mut^* NTOs stained for DAPI (blue) and FOXA2 (green) upon RA, RA+BMP4 (25 ng/ml), RA+NOG (200 ng/ml) and RA+LDN (500 nM). Scale bars, 100 µm. **E:** 3D analysis of proportion of FOXA2^+^ NTOs shown in B and D of WT (purple) and *Nog^mut^* (cyan). Results were statistically analysed using an ordinary one-way ANOVA, always comparing WT with *Nog^mut^* samples. RA vs RA *p* = <0.0001, RA+BMP4 (10 ng/ml) vs RA+BMP4 (10 ng/ml) *p* = 0.0004, RA+BMP4 (25 ng/ml) vs RA+BMP4 (25 ng/ml) *p* = 0.0039, RA+LDN (500 nM) vs RA+LDN (500 nM) *p* = 0.0229, while the other conditions were n.s. A total of *n* = 3407 (between 306 and 414 per condition) WT NTOs and *n* = 3425 (between 328 and 415 per condition) *Nog^mut^* NTOs were included in the analysis. Concentration [x] expressed in ng/ml for BMP4 and NOGGIN, or in nM for LDN. Asterisks indicate statistical significance, n.s. indicates non-significance. **F:** 3D analysis of average surface area of FOXA2^+^ clusters in WT (purple) and *Nog^mut^* (cyan) NTOs. Each dot represents an average cluster area in a given NTO. Results were statistically analysed using Kruskal-Wallis test and Dunn’s multiple comparisons test comparing RA with RA+BMP4 (in WT NTOs), RA and RA+NOG (in *Nog^mut^* NTOs). WT RA vs WT RA+BMP4 *p* = <0.0001, WT RA vs *Nog^mut^* RA *p* = <0.0001, and WT RA vs *Nog^mut^* RA+NOG was n.s. Asterisks indicate statistical significance, n.s. indicates non-significance. **G:** Representative images (single *z* slice) of a Day 6 WT NTO stained for DAPI (blue), FOXA2 (green), SHH (magenta) and NKX6.1 (cyan) for RA, RA+BMP4 (25 ng/ml), RA+NOG (200 ng/ml) and RA+LDN (500 nM). Scale bars, 100 µm. **H:** Representative images (single *z* slice) of a Day 6 *Nog^mut^*NTO stained for DAPI (blue), FOXA2 (green), SHH (magenta) and NKX6.1 (cyan) for RA, RA+BMP4 (25 ng/ml), RA+NOG (200 ng/ml) and RA+LDN (500 nM). Scale bars, 100 µm. **I:** Transcriptional response in FOXA2^+^ cells after BMP signal activation or suppression shows FOXA2^+^ cells induce *Noggin* expression after BMP exposure. Schematic: at Day 3, BMP perturbations were added for 8 hours before NTOs were dissociated and FACS sorted into FOXA2^+^ and FOXA2^-^ cells for RNA analysis. Expression levels (cpm) of *Noggin* in FOXA2^+^ cells upon RA, RA+BMP4 (1.5 ng/ml) and RA+LDN (100 nM) treatments. Results were statistically analysed using DESeq2 with pairwise comparison, RA+BMP4 vs RA *p* = <0.0001 for *Noggin* expression. Asterisks indicate statistical significance, n.s. indicates non-significance. **J:** Model of conceptional interactions between BMP4 and NOGGIN activities in FOXA2^+^ cells and their interaction via pSMAD 1/5/9 activity in FOXA2^-^ cells.

BMP4 ligand treatment led to a dose-dependent decrease in the proportion of NTOs with any FOXA2^+^ clusters persisting at Day 6 (Figure 4B-C). Conversely, BMP pathway inhibition by application of recombinant NOGGIN or the small molecule LDN193189 led to a mild increase in the total proportion of NTOs with any FOXA2^+^ clusters (Figure 4B-C, S3E). When broken down into whether each NTO contained 1, 2 or 3+ clusters, BMP inhibition resulted in a significant increase in the proportion of NTOs with 3+ endpoint clusters (Figure 4C). This is consistent with a role of endogenous BMP signalling in promoting FOXA2 cluster extinction, which is alleviated by BMP inhibition.

We then tested whether endogenously produced NOGGIN plays a role in regulating FOXA2^+^ clusters in NTOs by generating *Noggin* mutant (*Nog^mut^*) ESCs with CRISPR-Cas9 (Figure S4F), then subjecting them to NTO formation and BMP pathway perturbations (Figure 4D-F). A significantly lower proportion of *Nog^mut^* NTOs contained any endpoint FOXA2^+^ clusters compared to WT NTOs (Figure 4E). BMP4 treatment led to a further and dose-dependent reduction in the proportion of *Nog^mut^* NTOs with any FOXA2^+^ clusters (Figure 4E). Furthermore, when we quantified the size of FOXA2^+^ clusters, *Nog^mut^* NTOs and BMP-treated WT NTOs both exhibited significantly smaller FOXA2^+^ clusters than untreated WT NTOs (Figure 4F). Exogenous application of NOGGIN protein to *Nog^mut^*NTOs rescued the proportion of FOXA2^+^ NTOs (Figure 4E) and FOXA2^+^ cluster size (Figure 4F) to WT levels. We confirmed that the FOXA2 clusters in NOGGIN-treated *Nog^mut^* NTOs represent functional SHH-expressing floorplates at Day 6 by co-staining for SHH and SHH/GLI-responsive NKX6.1 (Figure 4G-H).

Together, these data demonstrate that BMP signalling activity level regulates FOXA2^+^ cluster outcomes during NTO self-organisation. BMP pathway activation versus inhibition controlled FOXA2^+^ cluster extinction versus persistence in a dose-dependent manner, as well as modulating FOXA2^+^ cluster size. Endogenously, BMP4 ligand and BMP-inhibitor NOGGIN were both expressed by FOXA2^+^ cells (Figure 3C), leading us to query the regulatory interactions between FOXA2, BMP4 and NOGGIN that underpin self-organisation in NTOs. To examine how BMP signalling affects gene expression response in FOXA2^+^ cells, we perturbed the BMP pathway in NTOs then sorted according to FOXA2-Venus reporter level and performed RNA-seq after 8 hours, in order to capture early transcriptional responses in nascent FOXA2^+^ cells (Figure 4I, S4G-I). FOXA2^+^ cells from BMP4-treated NTOs showed a significant increase in *Noggin* expression (Figure 4I), in agreement with BMP-driven *Noggin* induction reported in human 2D gastruloids ^21^, indicating a feedback loop in which FOXA2^+^ clusters protect themselves from the BMP ligands they themselves produce (Figure 3C).

In summary, we propose a FOXA2-BMP4-NOGGIN signalling circuitry that underpins cluster competition during NTO self-organisation (Figure 4J). FOXA2 cells express BMP ligands (Figure 3C) that, plausibly through pSMAD1/5/9, suppress FOXA2 in surrounding cells. NOGGIN, also expressed by FOXA2 cells (Figure 3C) and under feedback control by BMP (Figure 4J), promotes the persistence and size of “winner” clusters.

### NOGGIN regulates floorplate size in the anterior NT *in vivo*

Given our observations in NTOs we asked whether deleting *Noggin* might reduce floorplate size *in vivo*. We crossed mice bearing *Noggin* floxed (*Nog^Fl/Fl^* or *Nog^Fl/+^*) ^22^ and *Sox2^CreERT^*^2^*^/+^* ^23^ alleles (Figure 5A, S5A), tamoxifen-treated at E5.5 to induce *Nog* conditional knockout (cKO) in the epiblast and harvested embryos at E10.5 to evaluate FOXA2 and SHH expression in the NT at different anatomical levels (Figure 5B).

**Figure 5:**
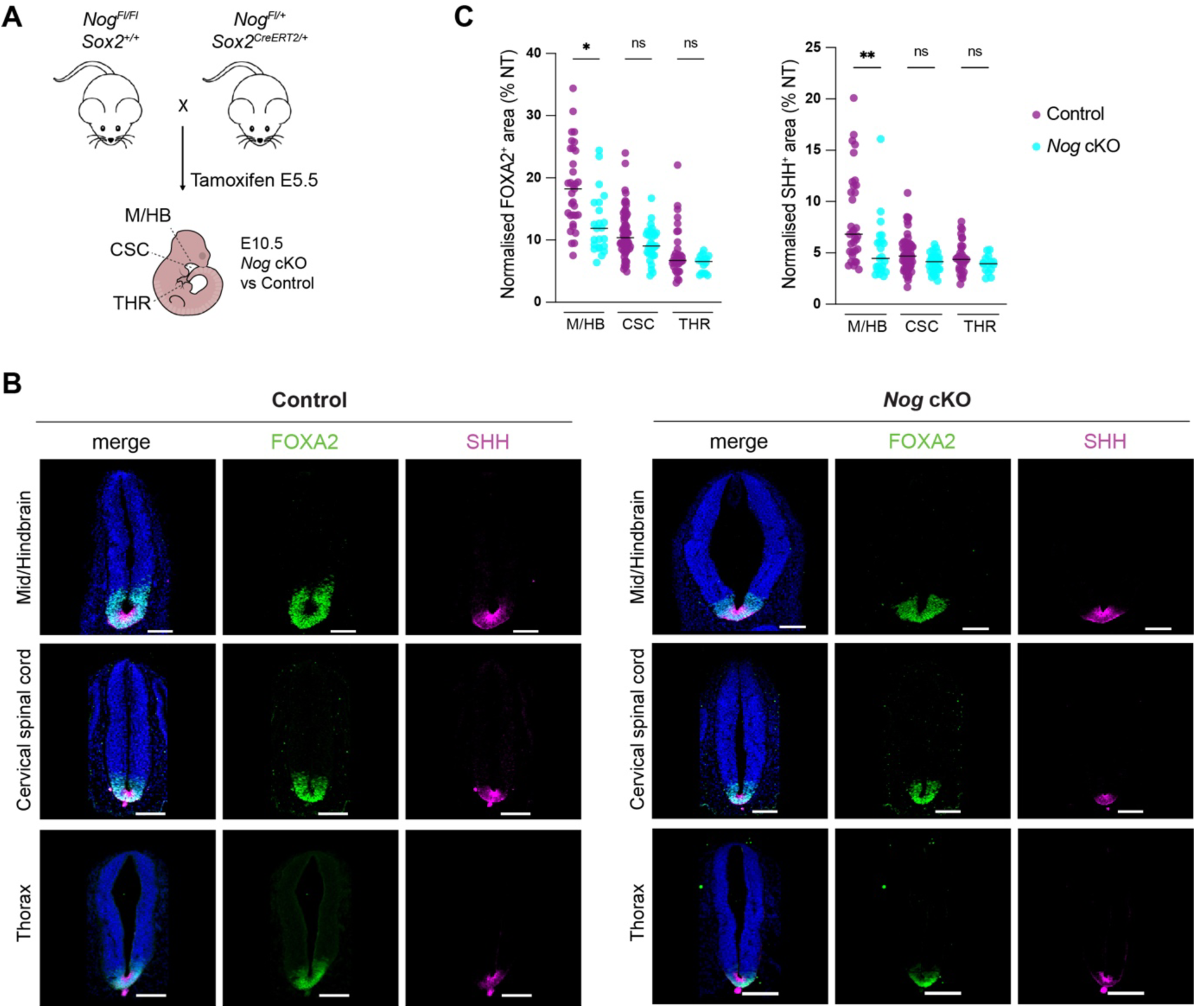
*In vivo* role of NOGGIN in anterior floorplate size in the mouse embryo. **A:** Schematic of mouse crossing performed for inducible deletion of *Noggin*. E10.5 mouse showing the anatomy and section planes of the harvested and analysed embryos. **B:** Representative images of E10.5 Control (*Nog^Fl/+^;Sox2^+/+^* or *Nog^Fl/Fl^;Sox2^+/+^*) and *Nog* cKO (*Nog^Fl/Fl^;Sox2^CreERT^*^2^*^/+^*) embryos of Mid/-Hindbrain (M/HB), Cervical spinal cord (CSC) and Thorax (THR) regions stained for DAPI (blue), FOXA2 (green) and SHH (magenta). Scale bars, 100µm. **C:** Normalised area of FOXA2 and SHH at different anterior-posterior levels of the embryonic NT. Quantification was made on serial sections of 6 Control embryos (*N* = 136 sections total, 33 for M/HB, 62 for CSC, 33 for THR) in purple and 3 *Nog* cKO embryos (*N* = 68 sections total, 23 for M/HB, 30 for CSC, 15 for THR) in cyan from two litters. Kruskal-Wallis test and Dunn’s multiple comparison tests were performed between anatomical regions of Control vs *Nog* cKO (FOXA2: M/HB vs. M/HB *p* = 0.0165, CSC vs. CSC *p* = 0.2706, THR vs. THR *p* = 0.3926, SHH: M/HB vs. M/HB *p* = 0.0031, CSC vs. CSC *p* = 0.0792, THR vs. THR *p* = 0.4484). Horizontal bars represent means of the data, asterisks indicate statistical significance, n.s. indicates non-significance.

By quantifying the DAPI^+^ area in cross-sections, we observed that mutant embryos exhibited slightly smaller NTs in the cervical spine region compared to control embryos from the same developmental stage (Figure S5B). To avoid any confounding effect, we measured the areas of FOXA2 and SHH expression (Figure S5C-D) then normalised to respective NT size (Figure 5C). We found a significant decrease in the normalised areas of FOXA2 and SHH expression in *Nog* cKO compared to control embryos specifically in the mid-hindbrain region, in which the floorplate forms prior to notochord development ^24^. By contrast, there was no difference in the FOXA2 and SHH domain sizes in the notochord-dependent cervical and thoracic spinal cord. Altogether, our *in vitro* and *in vivo* findings suggest that floorplate establishment in the mid-hindbrain region is regulated via BMP-dependent cell communication.

## Discussion

Here, we describe how a floorplate signalling centre self-organises in clonal 3D NTOs. Following a pulse of RA signal, FOXA2^+^ cells emerge in a salt-and-pepper fashion at Day 3, then undergo a process of cluster formation and resolution to form a localised domain, in which SHH morphogen expression commences from Day 5 (Figures 1-2). During NTO self-organisation, intermediate FOXA2^+^ clusters do not behave independently, but instead promote each other’s extinction (Figure 2) via a BMP-mediated signalling mechanism, whilst NOGGIN protects FOXA2 (Figures 3-4). Furthermore, BMP4 application or *Noggin* mutation decreases FOXA2 domain size, which can be rescued by exogenous NOGGIN (Figure 4). Strikingly, the expression of BMP ligands and inhibitors are both enriched in the FOXA2^+^ cells, whereas BMP signal transduction indicated by pSMAD1/5/9 is enriched in FOXA2^-^ cells (Figure 3). This represents a self-organisation mechanism in which spatially scattered FOXA2^+^ cells express the signalling machinery driving their competition to result in a ventral pole. Future advances will allow us to understand cluster competition in NTOs more deeply, such as how FOXA2 levels or cluster size are controlled and how these properties relate to the ultimate outcome. A thorough understanding will also involve developing the means to determine the localization and dynamics of the BMP pathway components.

Expression of both the FOXA2 “inhibitor” (BMP4) and “activator”/protector (NOGGIN) molecules from the same cells (FOXA2^+^), and the feedback by which BMP4 promotes *Noggin* expression in FOXA2^+^ cells (Figure 4), displays features of a reaction-diffusion (RD) mechanism ^25, 26^. In the future, it will be interesting to test the extent to which an RD-type mechanism underpins floorplate patterning in NTOs. This NTO system also offers the intriguing possibility to define quantitatively how the relative contributions of long-range communication versus physical cell sorting mechanisms (Figure 2) complement each other to promote self-organisational robustness. It will also be interesting to see how our findings from mouse stem cells compare to recently described human NTOs that induce full or partial DV patterning ^27–29^. In those studies, fixed timepoint analysis showed scattered FOXA2 prior to floorplate formation, but as less than 30% of human NTOs formed floorplates, it has been unclear whether the scattered starting point was an intermediate to the final state, or whether distinct subpopulations arose over time ^27^. WNT signalling has been proposed to act in an RD patterning system in human NTOs ^29^, which we did not observe in mouse NTOs. Whether these discrepancies are due to experimental approaches or whether they highlight intriguing inter-species differences remains to be addressed.

Other 3D organoid models that form from aggregates of 100s-1000s of cells ^30–34^ can self-organise by amplifying initial asymmetries, yielding persistence of a signalling centre at the location of its initial emergence. In contrast, our NTOs form clonally from single ESCs and show a regulative self-organisation process, with nascent clusters competing via long-range signalling prior to establishing a stable ventral organiser. To our knowledge, cluster competition has not been observed in other organoid systems, although few studies have yet interrogated the earliest stages of their self-organisation due to constraints in efficiency, topographic predictability or optical density that have limited tractability. Recent works addressing how 3D gastruloids initiate self-organisation of an anterior-posterior axis ^35, 36^ have combined improvements in homogeneity of input cells and analytical tools to elucidate how early variation in Nodal/BMP and cell position act prior to heterogeneity in WNT activity that determines which cells will form the posterior pole, reinforced by cell sorting. Endogenous and exogenous BMP signalling have been shown to mediate organised lineage induction in 3D models of human epiblast ^37, 38^ and mouse appendages ^39^, but again without evidence of cluster competition. The processes we describe here are also distinct from those found in clonal intestinal organoids where local activation of YAP1 induces symmetry breaking via DLL1 to determine the future crypt ^40^, without indication of competition between emergent DLL1 sites. An interesting question is whether mechanisms such as those we describe may be upstream of self-organisation mechanisms in other organoids.

Here we have identified and tested the importance of BMP signalling for cluster interactions in mouse NTOs. Since BMP is a key morphogen driving dorsal patterning of the NT from the dorsal roofplate organiser ^41–43^, it seems surprising to observe BMP ligand expression in ventral floorplate cells. However, BMP ligands are reported to be expressed in the *in vivo* floorplate ^44–46^, in keeping with the enrichment of BMP ligand expression we observe in FOXA2^+^ cells of NTOs (Figure 3). Our observation that BMP4 and NOGGIN regulate floorplate formation and size in NTOs led us to investigate whether BMP signalling circuitry plays a role in floorplate formation *in vivo*. Conditional knockout of *Noggin* in the epiblast led to a reduction in floorplate size at E10.5, particularly at mid/hindbrain regions of the NT where the floorplate, like our NTOs, forms in the absence of underlying notochord (Figure 5). This raises the possibility that ventral sources of BMP may also contribute to NT patterning defects in mouse mutants of the BMP pathway ^44–46^.

The broader applicability of the BMP-NOGGIN circuit regulating floorplate formation in NTOs highlights the potential of tractable *in vitro* self-organising systems to uncover mechanisms that may contribute *in vivo*, away from the complexity of the developing embryo. This raises fascinating possibilities for the future: by integrating classical developmental biology with self-organisation in stem cell models, we can quantitatively tease apart how embryonic context and inductive cues interface with self-organising modules, to allow regulative yet robust outcomes of vertebrate development.

**Extended Figure S1:**
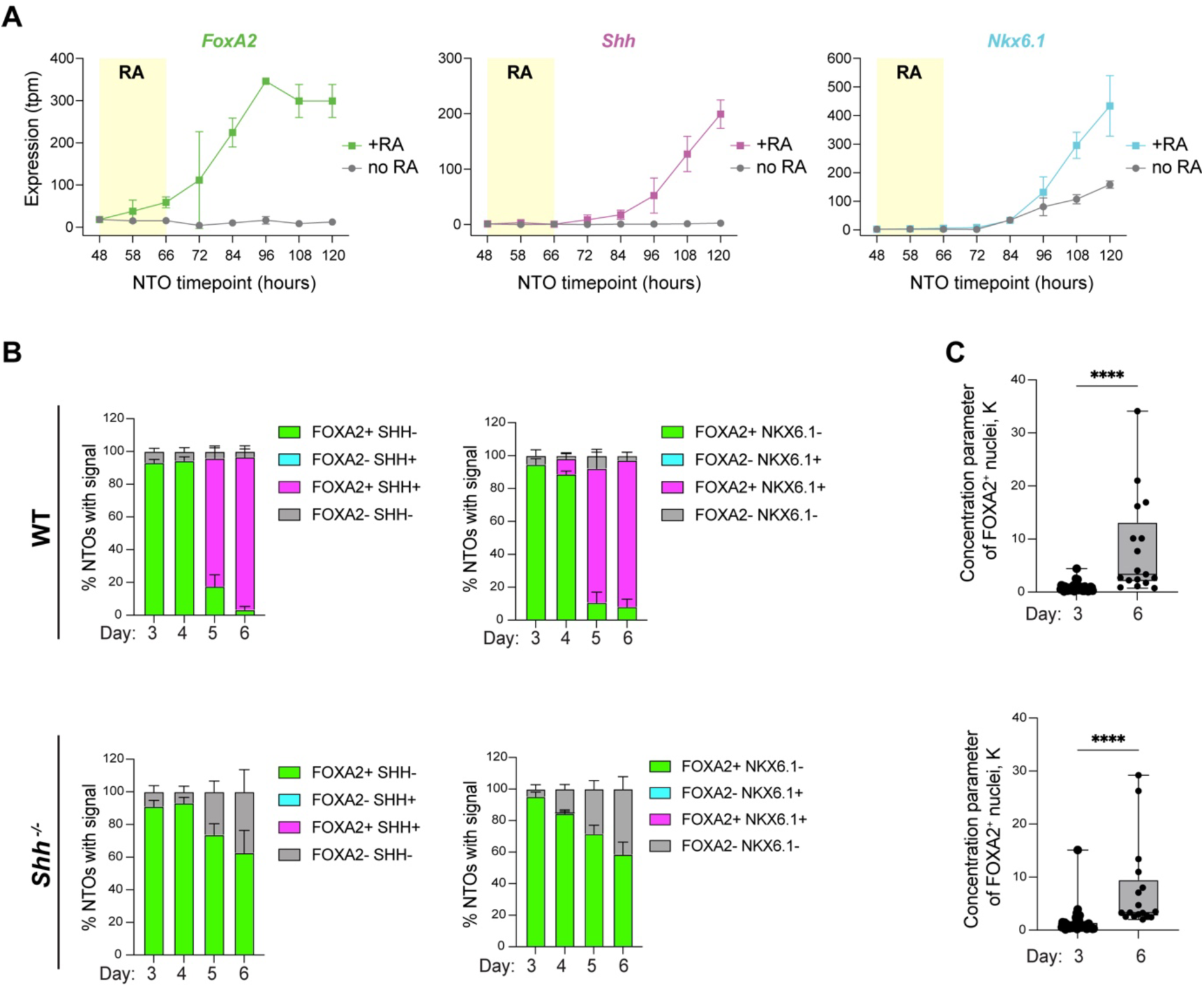
Characterisation of WT and *Shh^-/-^* NTO progression. **A:** Bulk RNA-seq of NTO timecourse shows expression levels of FOXA2, SHH and NKX6.1 matching the colours of Figure 1B in transcripts per million (tpm) ^19^. **B:** 3D quantification of marker expression in *Shh^-/-^* and parental WT NTOs over time with colours corresponding to *Shh^-/-^* staining in Figure 1D. *n* = 2011 NTOs for *Shh^-/-^* and *n* = 2239 NTOs for WT were used in the analysis of FOXA2/SHH, *n* = 1142 NTOs for *Shh^-/-^* and *n* = 1235 NTOs for WT were used in the analysis of FOXA2/NKX6.1. **C:** Concentration parameter, K, measurements of *Shh^-/-^* NTOs and the corresponding parental WT cell line. Each dot represents the K value of a single NTO (WT: *n* = 42 for Day 3, *n* = 17 for Day 6, *Shh^-/-^*: *n* = 41 for Day 3, *n* = 17 for Day 6). Results were statistically analysed using a Mann-Whitney test with *p* = <0.0001. Asterisks indicate statistical significance.

**Extended Figure S2:**
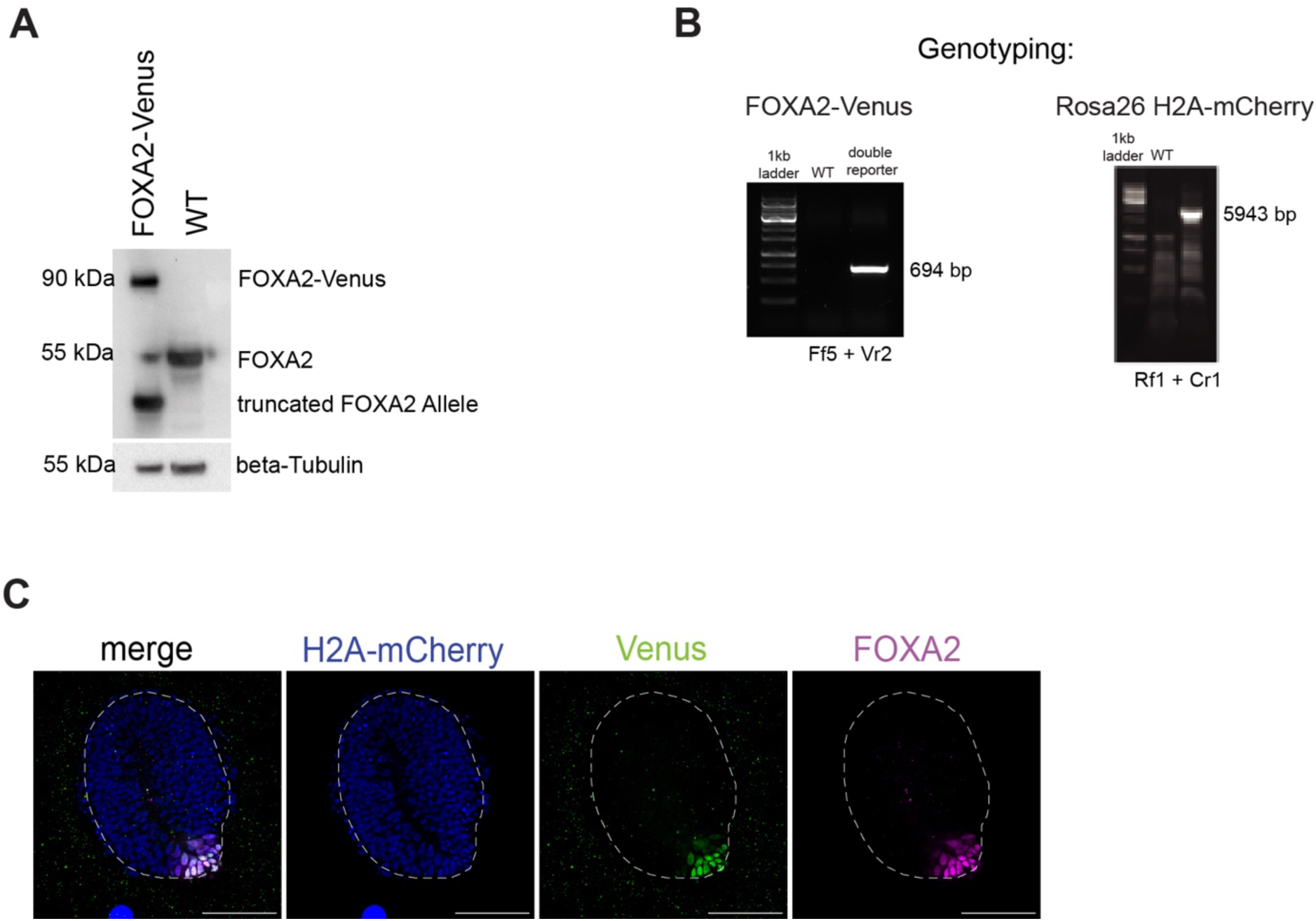
Characterisation of FoxA2-Venus knock-in line. **A:** Western Blot of FOXA2-Venus reporter line and WT detecting FOXA2 WT protein, FOXA2-Venus and a truncated FOXA2 allele. Beta-Tubulin served as a loading control. **B:** PCR confirmation of FOXA2-Venus and Rosa26 H2A-mCherry knock in. For Venus insertion, FoxA2_fwd5 (Ff5) and Venus_rev2 (Vr2) (product size: 694 bp), and for mCherry insertion, R26_leftout_fwd1 (Rf1) and mCherry_rev1 (Cr1) (product size: 5943 bp) were used. **C:** Representative Day 5 NTO (single *z* slice) showing co-expression of FOXA2-Venus and endogenous FOXA2 signal with H2A-mCherry (blue), Venus (green) and stained FOXA2 (magenta). Dashed lines mark the outlines of the NTO. Scale bars, 100 µm.

**Extended Figure S3:**
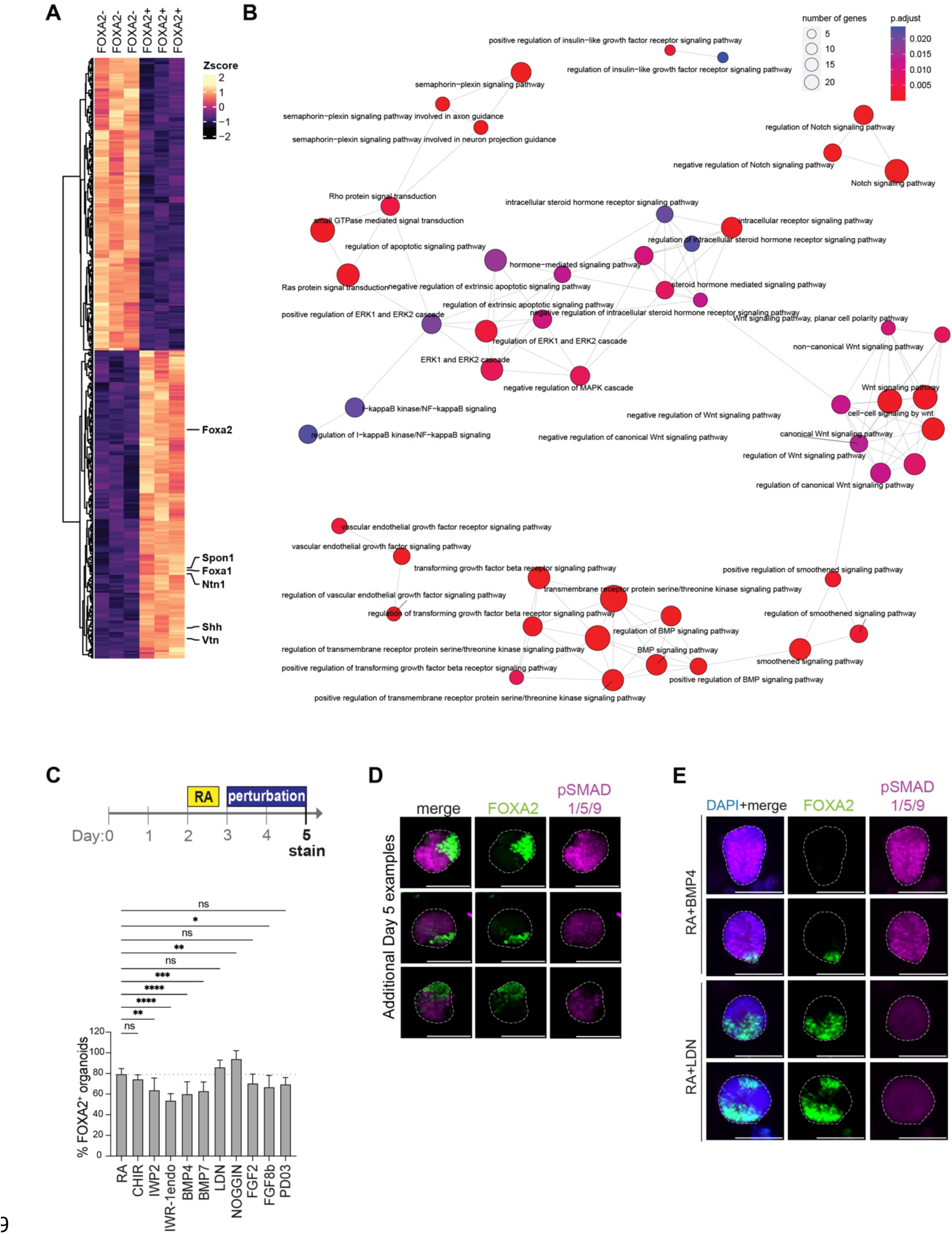
Analysis of RNA-seq data from FOXA2+ and FOXA2-cells and BMP analysis. **A:** Heatmap of clustering of top 500 differentially expressed genes from sorted Day 5 cells. **B:** Enrichment plots showing top 400 genes of various pathways from sorted Day 5 RNA-seq. **C:** Schematic of drug screen performed on Day 3-5 starting on Day 3, i.e. 6 hours post RA removal, with analysis of proportions of FOXA2^+^ NTOs on Day 5. The following concentrations were used: CHIR (3 µM), IWP2 (5 µM), IWR-1endo (1 µM), BMP4 (5 ng/ml), BMP7 (10 ng/ml), LDN (100 nM), NOGGIN (200 ng/ml), FGF2 (10 µg/ml), FGF8b (200 ng/ml), PD03 (1 µM). Results were statistically analysed using an ordinary one-way ANOVA and Dunnettes’s multiple comparisons test. RA vs RA+IWP2 *p* = 0.0019, RA vs RA+IWR-1endo *p* = <0.0001, RA vs RA+BMP4 *p* = <0.0001, RA vs RA+BMP7 *p* = 0.0009, RA vs RA+NOGGIN *p* = 0.0040, RA vs RA+FGF8b *p* = 0.0188, while all other tested conditions were n.s. Asterisks indicate statistical significance, n.s. indicates non-significance. **D:** Representative examples (single *z* slice) of Day 5 NTOs stained for FOXA2 (green) and pSMAD1/5/9 (magenta). Dashed lines mark the outlines of the NTOs. Scale bars, 50 µm. **E:** Representative examples (single *z* slice) of Day 5 NTOs stained for DAPI (blue), FOXA2 (green) pSMAD1/5/9 to visualise BMP pathway activity (right) (magenta) upon RA+BMP4 (1.5 ng/ml), and RA+LDN (100 nM). Dashed lines mark the outlines of the NTOs. Scale bars, 100 µm.

**Extended Figure S4:**
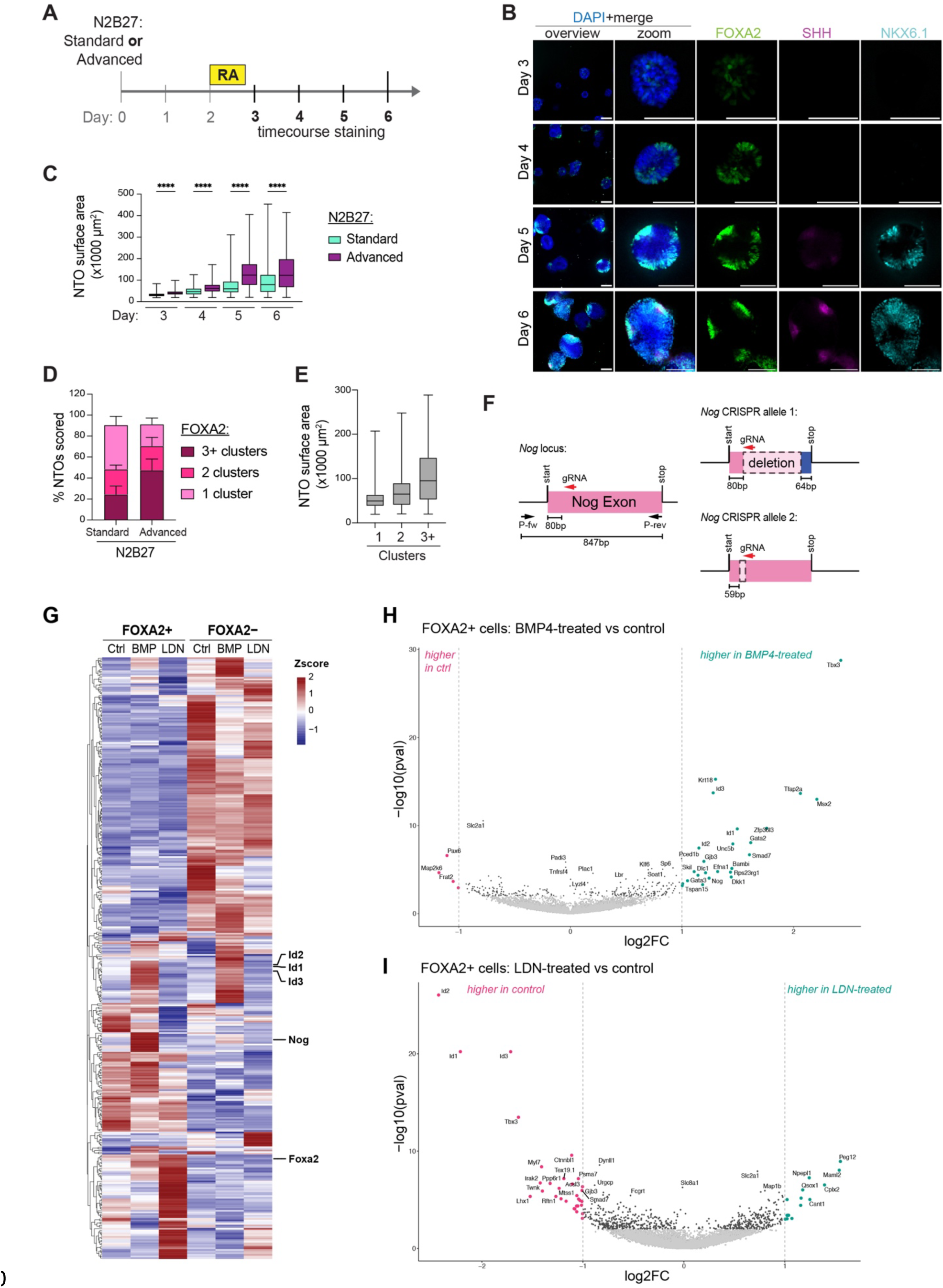
Characterisation of large NTOs and transcriptional induction data. **A:** Basic timeline of NTO production. **B:** IF timecourse of NTOs in rich medium (“Advanced”) fixed from Day 3-6 showing (from left to right, single *z* slice) a merged overview image at 10x magnification, and a representative NTO (zoom) stained for DAPI (blue) and the DV patterning markers FOXA2 (green), SHH (magenta), NKX6.1 (cyan). Scale bars, 100 µm. **C:** 3D analysis of NTO surface area in normal NTOs (standard medium, cyan, *n* = 2463) and large NTOs (advanced medium, purple, *n* = 1880) from data in Figure 1B and Figure S4B. Asterisks indicate statistical significance. **D:** 3D analysis of numbers of FOXA2^+^ clusters in WT NTOs on Day 6 from data in Figure 1B and Figure S4B (*n* = 751 Standard, *n* = 384 Advanced). **E:** 3D analysis of NTO surface area with 1, 2 or 3+ FOXA2^+^ clusters (advanced medium, total *n* = 1321 NTOs, *n* = 664 for 1 cluster, *n* = 284 for 2 clusters, *n* = 373 for 3+ clusters). **F:** Schematic of *Nog^mut^* cell line generation. The gRNA binds 80bp downstream of the start codon. Sequencing of the mutant clone confirmed a trans-heterozygous mutation. CRISPR allele 1: large deletion followed by a frameshift and premature stop codon, such that only 27 amino acids of NOGGIN protein remain, which correspond only to the signal peptide, followed by 21 out-of-frame amino acids. CRISPR allele 2: 33bp deletion including 8 amino acids of the signal peptide and 3 amino acids of NOGGIN. **G:** Heatmap showing clustering of top 200 differentially expressed genes from RNA sequencing. **H:** Volcano plot of genes showing Log2 fold change of expression in RA+BMP4 vs RA in FOXA2^+^ cells from RNA sequencing. Dark grey: *p*<0.01, Green (#009988): *p*<0.01 and FC>2, Pink (#EE3377): *p*<0.01 and FC<-2. **I:** Volcano plot of genes showing Log2 fold change of expression in RA+LDN vs RA in FOXA2^+^ cells from RNA sequencing. Dark grey: *p*<0.01, Green (#009988): *p* <0.01 and FC>2, Pink (#EE3377): *p*<0.01 and FC<-2.

**Extended Figure S5:**
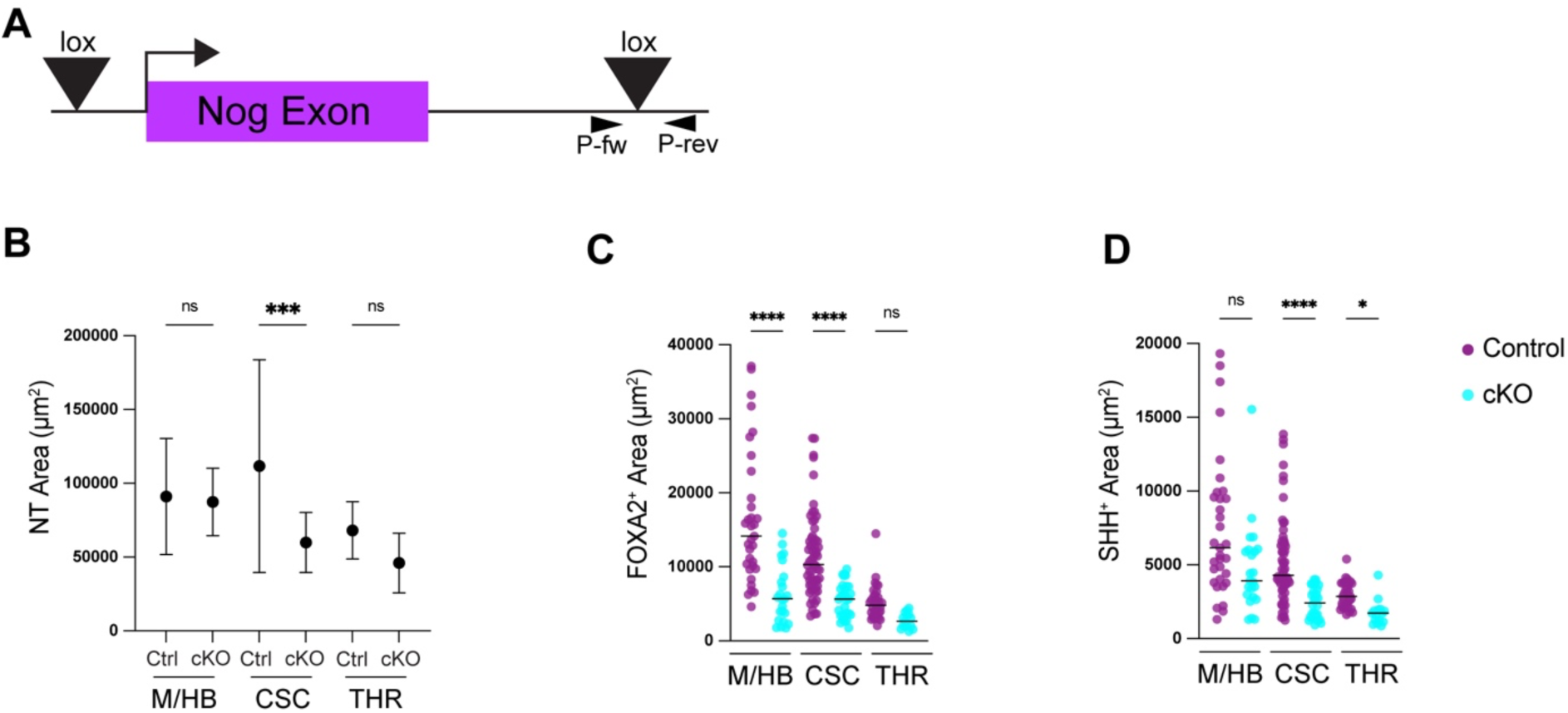
Characterisation of *Nog* cKO mouse. **A:** Schematic of *Nog* flox allele and the genotyping primers. **B:** Total area of the NT measured by DAPI signal in serial sections from the different A-P levels in of E10.5 Control (*Nog^Fl/+^;Sox2^+/+^*or *Nog^Fl/Fl^;Sox2^+/+^*) and *Nog* cKO (*Nog^Fl/Fl^;Sox2^CreERT^*^2^*^/+^*) embryos of Mid/-Hindbrain (M/HB), Cervical spinal cord (CSC) and Thorax (THR) embryos used to normalise the data in Figure 5C. The data is represented as mean ± SD. **C:** Raw area measurement of FOXA2 in sections from the different A-P levels corresponding to analysis from Figure 5B and C. Kruskal-Wallis test and Dunn’s multiple comparison tests were performed between anatomical regions of Control vs *Nog* cKO (M/HB vs. M/HB *p* = <0.0001, CSC vs. CSC *p* = <0.0001, THR vs. THR *p* = 0.0708). **D:** Raw area measurement of SHH in different anatomical regions from the different A-P levels corresponding to analysis from Figure 5B and C. Kruskal-Wallis test and Dunn’s multiple comparison tests were performed between anatomical regions of Control vs *Nog* cKO (M/HB vs. M/HB *p* = 0.1129, CSC vs. CSC *p* = <0.0001, THR vs. THR *p* = 0.0431).

## Materials & Methods

### Cell culture

#### Maintenance

Mouse embryonic stem cells (ESC) were cultured in N2B27+2i+LIF (2iLIF) and incubated at 37°C in 5% CO_2_. N2B27 medium comprised 1:1 DMEM/F-12 and Neurobasal (Gibco), 0.5% v/v N2 (Gibco), 1% v/v B27 (Gibco), 2 mM L-glutamine (Gibco), 1% penicillin/ streptomycin (Sigma), and 0.1 mM β-mercaptoethanol. To obtain 2iLIF medium, N2B27 was supplemented with 3 µM CHIR99021 (Tocris), 1 µM PD0325901 (Tocris) and 20 ng/ml murine LIF (Qkine). Tissue-culture treated 6-well plates were pre-coated with 0.15% gelatin (Sigma) in PBS at room temperature. Mouse ESCs were seeded at a density of 30,000 cells per 6-well, media was exchanged daily, and cells were passaged every third day by incubation with Accutase (Gibco) for 3 mins at 37°C, dissociation into single cells, centrifugation of the required number at 1500rpm for 5 mins, then re-seeding in a fresh well. R1 ESC line was used for most experiments and as the parental line for FOXA2-Venus reporter generation and for *Noggin* mutant (*Nog^mut^*). HM1 ESC background was used as the parental line for *Shh* knock-out (*Shh^-/-^*). Cell lines were routinely tested and confirmed negative for mycoplasma.

#### Generation of reporter cell lines

##### Generation of FOXA2-Venus reporter ESCs

Foxa2-Venus Fusion (FVF) E-85 plasmid (kind gift from Heiko Lickert ^47^) was used as a template for amplification of the FoxA2_exon3-venus-neo-PGK cassette, which was subsequently cloned into pGEMT-easy vector using Gibson Assembly technique. Two fragments for Gibson assembly were amplified using the E-085 vector as a PCR template:

1. Fragment1 - “FoxA2_exon3-Venus-Neo”:

GibAs_p3_fwd: CGGCCGCGGGAATTCGATAAAGAATCAAAGACCAGTGGA GibAs_p4_rev: GCCTCTCATTTCTACCGGGTAGGGGAGGCGCTTTT

1. 2) Fragment 2 - “3’ UTR-genomic DNA”:

GibAs_p1_fwd: TCCCCTACCCGGTAGAAATGAGAGGCTGAGTGGAGACTTT

GibAs_p2_rev2: CGCGAATTCACTAGTGATTCCCTCCTCCTTCAATTTCTCTTCCTTGTGTT

For CRISPR/Cas9 mediated knock-in of the Venus sequence at the 3’ end of the FoxA2 gene locus, gRNA3 (GCCTGCTAGCTCTGGTCACTG) was cloned into the pSpCas9(BB)-2A-Puro (PX459) vector, containing the Cas9 sequence and a gRNA expression cassette. For FoxA2 locus editing, both vectors were electroporated together into the ESC cell line R1 using the Amaxa Nucleofector 4D, program CG-104. 24 hours after the electroporation, G418 (100 μg/ml) was added to the culture medium to select for positive clones. Single clones were expanded, and correct integration of the Venus cassette was validated by PCR, sequencing and standard Western blot analysis.

In brief, protein of cell lysates was obtained by directly adding 5xSample Buffer and boiling samples at 95°C for 10 mins. The protein was loaded onto a NuPAGE^TM^ 4-12% Bis-Tris gel (Invitrogen) and run for 20 mins at 100V, then 90 mins at 150V, then proteins were blotted onto a PROTRAN nitrocellulose transfer membrane (Whatman) by TE 77 Semi-Dry Transfer Unit (Amersham). The membrane was blocked in 5% BSA in 1x Transfer Buffer (10x TB, MeOH, H_2_O) supplemented with Tween-20. Primary antibodies used: rabbit anti-FOXA2 (Cell Signalling Technologies, #8186S, 1:1000) and mouse anti-ß-tubulin (MPI-CBG antibody facility, 1:200).

FoxA2_fwd5: GACCTCAAGGCCTACGAACA

Venus_rev2: GTTGTGGCGGATCTTGAAGT (Product size: 694 bp)

##### Generation of FoxA2-Venus and H2A-mCherry double-reporter ESCs

H2A-mCherry was integrated into the Rosa26 safe harbour locus of FOXA2-Venus cell line. We first obtained a Rosa26 targeting vector with Puromycin selection cassette (pR26 CAG AsiSI/MluI, Addgene 74286, Ralf Kuehn lab, ^48^), linearized the vector with SwaI restriction digest, and used Gibson assembly to insert the Gateway destination sequence immediately 3’ to the splice acceptor site, and created a new vector termed pR26-sA-DEST. Subsequently, we performed a Gateway LR reaction between pR26-sA-DEST and pME-H2A-mCherry to create pR26-sA-H2-mCherry. To edit the Rosa26 locus, we identified a new high efficiency guide RNA sequence ACTGGAGTTGCAGATCACGA (Rosa26-gRNA1), and co-transfected p-Rosa26-gRNA1 and pR26-sA-H2-mCherry to FoxA2-Venus cells with Amaxa nucleofection. After 24 hours, cells were selected with 1 µg/ml Puromycin and individual clones were expanded. Correct integration was confirmed by Sanger sequencing and the length of the PCR amplicon from two primers:

R26_leftout_fwd1: GCTTGGTGCGTTTGCGGGGAT

mCherry_rev1: TTGGTCACCTTCAGCTTGGCG (Product size: 5943 bp)

#### Generation of knockout mouse ESC lines by CRISPR/Cas9

gRNAs were designed using the following tools: https://portals.broadinstitute.org/gppx/crispick/public or https://www.vbc-score.org/

##### Shh^-/-^

A short guide RNA (sgRNA) sequence (GCTGCTGGTGTGCCCCGGGCTGG) was cloned into pSpCas9(BB)-2A-Puro (PX459) V2.0 (Addgene plasmid #62988) as outlined in ^49^. 2 µg of PX459 V2.0 containing the sgRNA sequence were electroporated into 2×10^6^ DVI2 ESCs (HM1 background) using program A023 of the Amaxa Nucleofector II (Lonza) to ablate endogenous *Shh* gene expression. Electroporated cells were seeded onto a gelatin coated 10cm CellBind plate (Corning) and cultured in 2iLIF. The following day cells were treated with 1.5 µg/mL Puromycin (Sigma) for 48 hours. Cells were grown in 2iLIF until resistant colonies were ready to be pick. Single and well spaced-out colonies were picked using a 20-μl pipette tip, dissociated in 0.25% Trypsin (Gibco) and plated onto feeder cells in serum-containing ESC medium + LIF in a 96-well plate to allow expansion. Colonies were screened and confirmed for deletions via Sanger sequencing using primers ShhKOpFW (CAAGCTCTCCAGCCTTGCTA) and ShhKOpRV (CTGCTCCCGTGTTTTCCTCA). The 11bp deletion caused a frameshift and premature stop codon. Cell lines were adapted back to 2iLIF conditions.

##### Nog^mut^

sgRNA2 (GGCGGATGTGTAGATAGTGCTGG) was cloned into the U6-IT-EF1As-Cas9-P2A-GFP vector, containing the Cas9 sequence and a gRNA expression cassette. As described previously, ESCs (R1) were electroporated with the construct using an Amaxa Nucleofector 4D (program CG-104). 40 hours post electroporation, single GFP^+^ cells were analysed and sorted based on their GFP expression relative to non-fluorescent control on a FACS Aria III machine into individual wells of 96-well plates and clonal lines were expanded. One clone was confirmed as trans-heterozygous mutant with a different sized deletion in each allele by PCR and sanger sequencing using the primers Nog_fwd1 TGAGGTGCACAGACTTGGAT and Nog_rev2 GCCGCCTTCCCAAGTAGA. CRISPR allele 1: large deletion followed by a frameshift and premature stop codon. 27 amino acids of NOGGIN protein remain, only corresponding to the signal peptide, followed by 21 out-of-frame amino acids. CRISPR allele 2: 33bp deletion including 8 amino acids of the signal peptide and 3 amino acids of NOGGIN.

#### Neural tube organoid (NTO) formation

ESCs were dissociated into single cells as described in “Maintenance”. After centrifugation of the required number of ESCs in N2B27, the pellet was placed on ice then resuspended in Matrigel (Corning) to yield a suspension at 100 cells per µl of Matrigel. 45 µl of this suspension was spread evenly across a MatTek dish for FACS and RNA-seq experiments, or 4 µl of suspension was carefully distributed in the centre of an optical-bottom 96-well (PerkinElmer) for immunofluorescence experiments. Gellification was performed at 37°C for 15 mins (MatTek) or 4 min (96-well), then covered with pre-warmed N2B27 medium (2 ml per MatTek, or 100 µl per 96-well. “Standard” N2B27 medium for NTO generation was the same as the N2B27 base for 2iLIF (see “Maintenance”), with the addition of 1% sodium pyruvate (Gibco). “Advanced” N2B27 for making larger NTOs used Advanced DMEM/-12 (Gibco 12634-010 instead of 21331-020) and was supplemented with 1% v/v non-essential amino acids (Gibco). In both cases, medium was exchanged daily from Day 2 onwards. On Day 2, 250 nM all-trans RA (Sigma) was added to the N2B27 medium, then removed after 18 hours. For perturbations to NTOs, the following proteins or small molecules were added at the concentrations indicated in the corresponding figures: BMP4 (Gibco), BMP7 (Gibco), LDN (Selleck Chemicals), NOGGIN (R&D), CHIR99021 (Tocris), IWP2 (Selleck Chemicals), IWR-1endo (Sigma), FGF2 (PeproTech), FGF8b (in house), or PD0325901 (Tocris).

### Immunofluorescence

#### Wholemount staining of NTOs

NTOs were fixed in 2% PFA for 30 min at room temperature, by adding 1:1 volume of room temperature 4% PFA on top of culture media, to avoid Matrigel disintegration. After fixation, samples were washed three times in PBS, then blocked and permeabilised in b/p buffer (PBS, 1% BSA, 0.5% Triton X-100) for 24 hours at 4°C. Primary antibodies were applied in b/p buffer for 24-72 hours at 4°C to ensure deep penetration of the samples: LEF1 (Cell Signalling Technologies, #2230S, 1:200), FOXA2 (R&D, #AF2400, 1:400), FOXA2 (Cell Signalling Technologies, #8186S, 1:1000), NKX6.1 (DSHB, #F55A10s, 1:200). Samples were washed three times and for a total of at least 24 hours in b/p at 4°C, then incubated with Alexa Fluor conjugated secondary antibodies (Invitrogen Donkey anti-Mouse/Goat/Rabbit 1:400) and DAPI (Sigma) for 24-72 hours at 4°C. Finally, samples were washed then stored in PBS.

The following primary antibodies were also used with modifications to the fixation and staining procedures. For SHH (Cell Signalling Technologies, #2207S, 1:300), antigen retrieval with 1x citrate buffer (DAKO) was performed for 45 mins at 65°C prior to blocking and permeabilization. For pSMAD1/5/9 (Cell Signalling Technologies, #13820S, 1:800), fixation was performed in EMgrade PFA (Electron Microscopy Sciences) (8% diluted to a 2% final concentration) (12mins for 4µl Matrigel at room temperature). For both pSMAD1/5/9 and pERK (Cell Signalling Technologies, #4370S, 1:500), after fixation samples were permeabilised with methanol for 10 mins at -20°C, then blocked in db/p (PBS, 5% donkey serum, 0.3% Triton X-100) at 4°C.

#### Optical clearing of wholemount NTOs

PBS was removed and CUBIC-R+(N) refractive index matching solution ^50^ was added to completely cover the samples and incubated for 30-60 min at RT. After this adaptation, CUBIC-R+(N) solution was exchanged to fresh CUBIC-R+(N) supplemented with 2 mg/ml propylgallate (Sigma) to reduce photobleaching.

#### Staining of embryo sections

E10.5 embryos were fixed on ice for 75 mins in 4% PFA, washed in PBS, then adapted to sucrose and embedded in gelatin prior to cryosectioning. Slides were washed 3x for 15 mins each in 41°C PBS to dissolve the gelatin. Afterwards, slides underwent antigen retrieval as per SHH staining protocol above, then were blocked for 1 hour at room temperature in b/p. Primary antibodies as above were incubated overnight at 4°C in a humidified chamber. Samples were washed 3x for 5 mins with PBS+0.3% Triton X-100 at room temperature. Secondary antibodies were applied in a concentration of 1:200 in b/p and incubated overnight at 4°C. Samples were washed 3x for 5 mins with PBS+0.3% Triton X-100 at room temperature, then mounted in ProLong Diamond Antifade Mountant.

### Imaging

#### Viventis lightsheet live-imaging

ESCs were dissociated and seeded as described in “Neural tube organoid (NTO) formation”, but at a density of 80 cells/µl Matrigel in special Viventis chambers, lined with FEP film from inside. Chambers were manufactured manually by gluing the FEP sheet to the plastic chambers and left to dry overnight. Chambers were cleaned with plasma to reduce hydrophobic properties and facilitate the attachment of the Matrigel. Before use, chambers were washed 3x with water and 80% ethanol, and dried and sterilised under UV light for at 1 hour at room temperature.

After seeding and gellification of 40 µl Matrigel, 1.5 ml of N2B27 medium was added to the chamber and exchanged daily from Day 2. Live-imaging was initiated from Day 3 of the NTO generation procedure, i.e. 6 hours after RA removal, over a timecourse of 48-75 hours. Samples were incubated at 37°C in 5% CO2 with humidification throughout. Images were acquired using two Nikon 10x objectives (NA 0.3) for illumination and for detection one Nikon 25x (NA 1.1) water immersion objective with 18x magnification and imaged with the following settings: mCherry 50 ms exposure time, 0.5% laser power; Venus 10 ms exposure time, 3.5% laser power. Multiple NTOs at different *xyz* positions were imaged at each timepoint. Image acquisition was performed in stacks with 3 µm step size every 15 mins.

#### Spinning d isk fixed sample imaging

Wholemount NTOs and embryo sections were imaged on a Spinning Disk Confocal Olympus (inverted) microscope using a 10x (NA 0.4) air objective. Lasers were used at 100% and exposure time was adjusted to 200 ms for all lasers. Resolution was 0.65 μm in *x* and *y*. For wholemount NTOs, 125 planes with 2 μm spacing in *z* were acquired to allow for 3D reconstruction, and an automated stage was used for high-throughput acquisition with unbiased selection of multiple positions per 96-well using a grid. For embryo sections, planes with 2 μm spacing in *z* were acquired. For von Mises-Fisher analyses, fixed NTO images were acquired on the same microscope but using a 40x (NA 1.25) silicon oil objective, with 0.57 μm spacing in *z*.

### Image processing and quantification

The transition from the scattered FOXA2^+^ state to the intermediate clustered state was scored manually in Figures 1D and 2C-F. The transition is heterogeneous from NTO to NTO, and the tissue contains a high density of nuclei, so it was difficult to apply standardised parameters for automated image analysis. During manual analysis, a cluster was defined as at least three cells that were grouped together and displayed nuclear signal intensity of a minimum of 15% of Day 5 FOXA2 signal intensity.

Automated analyses were performed for Figures 1C, 4C, 4E-F, S1B, S3C and S4C-E as described below.

#### Analysis of Viventis movies

As mentioned above, manual classification was performed to analyse data in Figure 2C-F. to avoid biases, two people analysed two different datasets and confirmed findings from two different microscope methods with the same conclusion.

For intensity quantification of clusters over time in Figure 2D (right panel), we used maximum intensity *z*-projections, performed intensity-based thresholding for segmentation, and quantified the total FOXA2^+^ intensity for each cluster for individual frames with a Python script using scikit-image library. The movies were manually inspected from an orthogonal view in the *x*, *z* plane to ensure that there were no clusters overlapping in the *z*-direction.

For presentation, representative maximum intensity *z*-projections were made in ImageJ, or movies were exported as .mp4 files.

#### Analysis of fixed wholemount NTO images

Raw .vsi files were processed in ImageJ. For presentation, single representative *z* slices were exported as .tif files with set display ranges for each channel and without any binning in *x-y*.

##### von Mises-Fisher analysis

Imaris software was used to 3D reconstruct and visualise each NTO. FOXA2^+^ nuclei above a set intensity threshold were identified using the spots function in Imaris and applied as a batch script. The precision of automated FOXA2^+^ nucleus identification was checked and corrected manually. The geometric centre of the inscribed spot was considered the geometric centre of each nucleus and their *x, y*, *z* coordinates were used to calculate the concentration parameter, *k*, as the resulting vector calculated as a sum of all vectors built from the centre of the NTO to the centre of each FOXA2^+^ nucleus and normalised to the number of FOXA2^+^ nuclei.

##### Correlation analysis of FOXA2 vs LEF1, pERK, pSMAD1/5/9 signal intensities in NTOs

To analyse the spatial correlation of FOXA2 and signalling pathways, we used images of fixed NTOs prepared and acquired on the spinning disc microscope as described above. For each NTO, we used ImageJ to visually determine the midsagittal plane, draw a curved path with linewidth of 10 pixel within the tissue, measured the intensities of different channels along this path, and exported the results. Then, we used a Python script to calculate the Pearson correlation between the FOXA2 channel and a second channel representing LEF1, pERK, or pSMAD1/5/9. In this way, we obtained a single Pearson correlation per NTO.

##### 3D quantification of NTO features

For 3D image analysis, we first processed the raw .vsi files to isotropic .tif files using CLIJ ^51^. GPU-accelerated image processing library in ImageJ. Due to the “fish-bowl” effect of CUBIC-R+(N)’s high refractive index (RI = 1.5), the effective *z* steps were 3 μm. We therefore binned images in *x-y* from to 0.65 μm to 3 μm so that the 3D voxel dimensions were comparable (3 μm × 3 μm × 3 μm in the *x*, *y* and *z* directions).

To quantify FOXA2^+^ cluster number per NTO in 3D (Figures 4C, 4E, S4D-E), we used CellProfiler 4 ^52^. First, we segmented the NTO outer boundary by applying Otsu thresholding to the *z* stacks from the DAPI channel. Then, we segmented FOXA2^+^ clusters by applying a set intensity threshold to the *z* stacks from the FOXA2 channel and assigned these “child” clusters to their “parent” NTO. The FOXA2 intensity threshold was determined by comparison between positive control RA-treated versus negative control untreated endpoint NTOs and applied equally across all samples from the same experiment.

For scoring whether NTOs expressed any FOXA2, NKX6.1 and/or SHH (Figure 1C, S1B, S3C), the mean intensity of the corresponding channel was measured inside the DAPI segmented volume. To measure the size of NTOs (Figure S4C, S4E), the NTO surface area was inferred from the DAPI segmented volume assuming spherical shape and to overcome confounding crenulation of the mask surface.

##### Measurement of FOXA2^+^ cluster size

The quantification of FOXA2^+^ cluster properties for fixed NTOs shown in Figure 4F was generated using a combination of discrete geometry and computer vision techniques. First, down-sampled multi-channel image volumes were processed in Ilastik ^53^ to produce a high contrast segmentation of NTOs from background. Surface reconstruction methods contained in the TubULAR package ^54^ were then applied to these processed image volumes in order to build mesh triangulations approximating the epithelial mid-surface of each NTO. FOXA2 signal intensity on each mesh surface was quantified by sampling the image volume at subpixel resolution onto mesh triangles and then using angle-weighted averaging to compute a single intensity value for each mesh vertex.

FOXA2^+^ clusters were identified on a per-vertex basis using a custom consensus clustering algorithm, which produces a single “best” clustering by comparing results across an ensemble of input clusterings. Prior to clustering, the FOXA2 intensity on each vertex was normalised between [0,1] by a linear re-scaling between the 5th and 95th percentiles of all raw intensities measured on WT NTO given RA treatment. A single clustering in the ensemble was generated by assigning an index to each vertex in the surface mesh using a modified k-means method. Given a set of random initial cluster centres, a modified version of Lloyd’s algorithm was used to greedily minimise the following cost function:

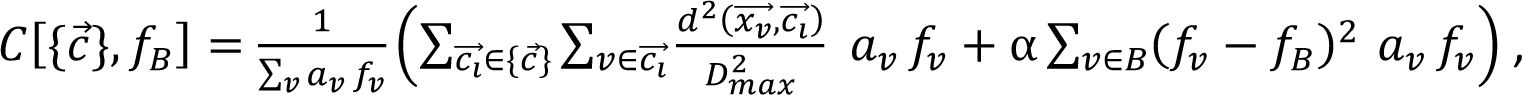

where {*c⃗*} denotes the set of all cluster centers, 𝑓_*B*_ is the average FOXA2 intensity of the background, 𝑎_/_is the barycentric area of vertex 𝑣, 𝑓_/_ is the normalised FOXA2 intensity on vertex 𝑣, *d*(*x⃗*, *y⃗*) is the 3D location of vertex 𝑣, 𝑑(*x⃗*, *y⃗*) denotes the 3D Euclidean distance between two points *x⃗* and *y⃗* on the surface, 𝐷_4$(_is the maximum distance between any two points on the NTO’s surface, and α is a positive scalar that sets the relative weight of the foreground versus background terms in the cost function. The double sum in the first term runs over all vertices 𝑣 associated to each cluster centre (*c⃗_l_* ϵ {*c⃗*}. The sum in the second term runs over all vertices assigned to the background. A vertex 𝑣 was assigned to cluster *c⃗_l_* if

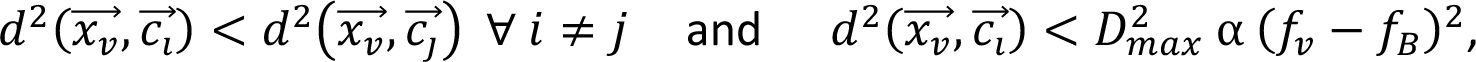

or alternatively to the background if

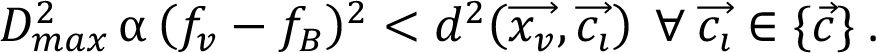

Intuitively, the cost function 𝐶[{*c⃗*}, 𝑓_*B*_] is minimised when all points in the background have similar (low) intensities and all foreground points are assigned to a nearby cluster centre. Given an ensemble of such clusterings, the method of ^55^ was used to produce a consensus clustering that is as close as possible to all the individual clusterings in the ensemble (formally it is the Fréchet sample mean of the clustering ensemble). The consensus clustering was then post-processed to ensure that each FOXA2^+^ cluster was simply connected. The use of a consensus clustering algorithm obviates the dependence of the output on the initial locations of the randomly chosen cluster centres. For our analysis, we set α = 1 and used 10 initial cluster centres to generate each element of the clustering ensemble. The initial background intensity was always taken to the lowest intensity on the surface. Numerical experiments showed that the choice of 𝑑(*c⃗*, *x⃗*) to be the 3D Euclidean distance produced indistinguishable results from more sophisticated methods using exact geodesic distances on the surface^56^ after consensus clustering. Total FOXA2^+^ surface area for a given NTO was taken to be the sum of all barycentric areas associated to each mesh vertex assigned to any FOXA2^+^ cluster, then the mean FOXA2^+^ surface area per cluster was computed based on the number of FOXA2^+^ clusters identified for each NTO.

##### Analysis of embryo sections

For area measurements of FOXA2 and SHH in embryo sections, we first converted raw .vsi files to maximum intensity z-projections in ImageJ. We set all channels to auto intensity and used the freehand selection tool from ImageJ to outline 1) the outer edge of the NT based on DAPI staining, 2) the NT lumen, 3) FOXA2 signal, and 4) SHH signal, and measured area in µm^2^. To get the actual area of the NT, we subtracted 1) NT area by 2) lumen area. To get the normalised area measurements of FOXA2 and SHH respectively, we divided the area measurement of the signal 3) or 4) by the NT area. For presentation, maximum intensity *z*-projections with set display ranges were used. We analysed 6 control and 3 cKO embryos from two litters.

### RNA sequencing

RNA-seq data generated in this study are available at the Gene Expression Omnibus (GEO) under accession codes GSExxxxxx (FOXA2^+^ vs FOXA2^-^ cells at Day 5, Figure 3) and GSExxxxxx (FOXA2^+^ vs FOXA2^-^ cells after BMP perturbations, Figure 4).

#### RNA-seq of RA-treated vs untreated timecourse from Day 2-5, Figure S1

Timecourse RNA-seq data ^19^ was obtained from GEO with accession code GSE214368. In brief, RNA-seq reads were trimmed using Trim Galore v0.5.0, filtered to remove abundant sequences using Bowtie 2 v2.3.4.1, aligned to the GRCm38 genome (Ensembl release 94) using STAR v2.6.0c and summarised per gene with featureCounts (subread v1.6.2). Further analysis was performed using DESeq2 v1.18.1.

#### RNA-seq of FOXA2+ vs FOXA2-cells at Day 5, Figure 3

RA-treated NTOs were extracted from Matrigel, dissociated into single cell suspension and FACS sorted into two cell populations based on FOXA2-Venus expression relative to non-fluorescent control on a FACS Aria III machine. For each sample, produced in three biological replicates, a minimum of 50,000 cells was sorted for subsequent RNA extraction. RNA was extracted from sorted cells using QIAGEN RNeasy Micro Kit and then underwent library preparation using the NEBNext Poly(A) mRNA Magnetic Isolation Module (E7490) and the NEBNext Ultra Directional RNA Library Prep Kit for Illumina (E7420), and bulk RNA sequencing on an Illumina NextSeq500 (single-end reads 75 bp).

RNA-seq reads were trimmed using Trim Galore v0.5.0, filtered to remove abundant sequences using Bowtie 2 v2.3.4.1, aligned to the GRCm38 genome (Ensembl release 94) using STAR v2.6.0c and summarised per gene with featureCounts (subread v1.6.2). Further analysis was performed using DESeq2 v1.18.1 and activated and inhibited upstream regulators were predicted using IPA Upstream Regulator Analysis based on the differentially expressed genes. Differentially expressed genes (DEG) of FOXA2^+^ versus FOXA2^-^ samples at a false discovery rate (FDR) threshold of 5% were subjected to IPA Upstream Regulator Analysis for predicting activated and inhibited upstream regulators. In addition, top 400 DEGs ranked by padj were used in gene set over-representation analysis with clusterprofiler 4.6.2 and GO.db 3.16.0.

#### RNA-seq of FOXA2^+^ vs FOXA2^-^ cells after BMP perturbations, Figure 4

NTOs were treated with RA at Day 2 for 18 hours as usual. At Day 3, i.e. 6 hours after RA removal, BMP4 (1.5 ng/ml) or LDN (100 nM) was added for 8 hours. To harvest cells, NTOs were washed twice with ice cold PBS, then dissociated in TryplE. FOXA2-Venus^+^ or FOXA2-Venus^-^ cells were FACS sorted into lysis buffer in a 96-well plate, with 100 cells sorted for each sample in triplicate, then snap frozen on dry ice. RNA was extracted and sequencing libraries were prepared following the TM3’seq protocol ^57^ for subsequent quantitative sequencing with enrichment for polyadenylated RNAs. Pooled libraries were sequenced on the Illumina NovaSeq S4 PE150 (paired-end 500bp reads).

The paired-end raw reads were first split and the 12bp of UMI were extracted from read2 with UMI-tools (v1.0.0). Reads were then trimmed with Cutadapt (v1.18) and mapped to the mouse genome (mm10) using STAR (v2.5.2a). UMI counts at gene levels were quantified by HTSeq (v0.11.2). The normalised counts per million (cpm) were obtained with DESeq2 (v1.26.0).

### Mouse procedures

Mouse procedures were approved under the license BMWFW-66.018/0006-WF/V/3b/2016 (“Coordination of tissue growth and patterning in neural tube development”) from the Austrian Bundesministerium für Wissenschaft, Forschung und Wirtschaft and performed in accordance with the relevant regulations. The following strains were previously described: *Sox2^CreERT^*^2^ ^23^, *Nog^Fl/+^* ^22^. To conditionally delete Noggin, *Nog^Fl/+^* were bred to *Sox2^CreERT^*^2^ mice to generate *Nog^Fl/Fl^*, *Sox2^CreERT^*^2^*^/+^* embryos. Pregnant mothers were intraperitoneally injected with 4mg tamoxifen in sunflower oil per mouse at embryonic Day E5.5. Embryos were harvested at E10.5. The expected ratios of genotypes were obtained.

#### Genotyping

Lysates for PCR were prepared following the HotSHOT protocol ^58^. Briefly, yolk sacs were transferred to clean tubes and lysed in Alkaline lysis reagent (25 mM NaOH, 0.2 mM disodium EDTA, pH 12) at 95°C for approx. 45 min. After cooling samples to 4°C, 1x volume of Neutralization buffer (40 mM Tris-HCl, pH5) was added to stop the lysis reaction. 2 µl of lysate was used in a 20 µl PCR reaction. To genotype the *Nog^FL^*, Primers Nog F 13351 (CCA CAA TAT CCA GCC CTT GT) and Nog R 13352 (AAG AGG CCC ATG TGA GTG TC) were used to detect the floxed allele. 10 µl PCR product was loaded onto a 2% gel and run at 110 Volt for 30 min. The WT band was 186bp, the mutant band 300bp. To genotype the *Sox2^CreERT^*^2^, primers 344 (GTCCAATTTACTGACCGTACACC) and 345 (GTTATTCGGATCATCAGCTACACC), as well as the internal control primers OIMR0042 (CTAGGCCACAGAATTGAAAGATCT) and OIMR0043 (GTAGGTGGAAATTCTAGCATCATCC) were used. The Mutant band was 705bp, the Internal Control 324bp.

## Data representation & availability

Data of this study are presented as follows. Bar-plots with error bars showing standard deviation (SD): Figures 1C, 1E, 2E, 4C, 4E, 4I, S1B, S3C, S4D. Box-plots with median represented as centre line, whiskers show range of values, dots (if present) represent all measurement points: Figures 1F, 3E, S1C, S4C, S4E. Scatter dot-plots with mean represented as centre line (or dot) and error bars showing standard deviation (SD): Figures 4F, 5C, S5B, S5C. Asterisks indicate statistical significance, n.s. indicates non-significance.

## Data availability

RNA-seq data generated in this study have been deposited at GEO under accession codes GSExxxxxx (FOXA2^+^ vs FOXA2^-^ cells at Day 5, Figure 3) and GSExxxxxx (FOXA2^+^ vs FOXA2^-^ cells after BMP perturbations, Figure 4).

Primers and oligonucleotide sequences are available in the Materials & Methods section.

## Supplementary files provided

**Supplementary Movies 1-2**

Live-imaging of RA-treated NTO showing FOXA2 self-organisation from Day 3-5. Maximum intensity *z*-projections of H2A-mCherry (magenta) and FOXA2-Venus (green) merged (Movie 1) or FOXA2-Venus (green) single channel (Movie 2) are represented as movies, related to Figure 2C.

**Supplementary Movies 3-4**

Live-imaging of RA-treated NTO showing FOXA2 self-organisation from Day 3-5. Maximum intensity *z*-projections of H2A-mCherry (magenta) and FOXA2-Venus (green) merged (Movie 3) or FOXA2-Venus (green) single channel (Movie 4) are represented as movies, related to Figure 2D, Extinction.

**Supplementary Movies 5-6**

Live-imaging of RA-treated NTO showing FOXA2 self-organisation from Day 3-5. Maximum intensity *z*-projections of H2A-mCherry (magenta) and FOXA2-Venus (green) merged (Movie 5) or FOXA2-Venus (green) single channel (Movie 6) are represented as movies, related to Figure 2D, Sorting.

## Author contributions

T.K., H.T.S., and E.G. designed and performed experiments, created methodologies, and analysed the data. K.I. provided supervision, set up live imaging conditions, developed image analysis pipelines and helped analysing data. D.C. and E.S. developed image analysis pipelines. M.M. and J.B. generated *Shh^-/-^* ESCs. J.W. performed bioinformatic analyses. E.C. and L.A. performed experiments. S.L. and A.K. provided *Nog* cKO embryos. A.H. and R.N. assisted with pharmacological screen. N.E. supported development of larger NTO media. H.T.S., T.K. and E.M.T. wrote the manuscript. E.M.T. conceived and supervised the study.

## Acknowledgements

We thank P. Pasierbek, A. C. Moreno, T. Lendl and K. Aumayr for microscopy support, G. Schmauss for FACS support, M. Novatchkova for assistance with Bioinformatic analyses, J. Ahel, S. Horer, E. Cesare, E. Norouzi for technical assistance, A. Meinhardt for supervision, DRESDEN-concept Genome Center, A. Vogt and the Vienna Biocentre NGS facility for RNA sequencing. We are grateful to M. Placzek and E. Martí for discussion about the floorplate, to S. Shvartsman for valuable input, to A. Aszodi, W. Masselink and S. Raiders for advice on statistical analyses, to J. Cornwall Scoones, G. Martello and Tanaka lab members for critical reading of the manuscript, E. Bassat and E. Chatzidaki for contributing schematics, and to K. Lust for support. The work was supported by funding from the Austrian Science Fund (FWF): F7803-B (Stem Cell Modulation) to E.M.T. and A.K, WWTF 10.47379/LS17037 and ERC AdG 742046 to E.M.T., Sir Henry Wellcome postdoctoral fellowship to H.T.S., ELBE fellowship to K.I., National Science Foundation (US) Phy 2013131 to E.S.. The A.K. lab is also supported by ISTA, the European Research Council under Horizon Europe grant 101044579, and S.L. is supported by Gesellschaft für Forschungsförderung Niederösterreich m.b.H. fellowship SC19-011. This work was supported in part by the Francis Crick Institute which receives its core funding from Cancer Research UK (CC001051), the UK Medical Research Council (CC001051), and the Wellcome Trust (CC001051). For the purpose of Open Access, the authors have applied a CC BY public copyright license to any Author Accepted Manuscript (AAM) version arising from this submission.

## References

1. Spemann, H. & Mangold, H. über Induktion von Embryonalanlagen durch Implantation artfremder Organisatoren. Arch. Für Mikrosk. Anat. Entwicklungsmechanik 100, 599–638 (1924).

2. Gurdon, J. B. Embryonic induction — molecular prospects. Development 99, (1987).

3. Martinez Arias, A. & Steventon, B. On the nature and function of organizers. Dev. Camb. Engl. 145, dev159525 (2018).

4. Sato, T. et al. Single Lgr5 stem cells build crypt-villus structures in vitro without a mesenchymal niche. Nature 459, 262–265 (2009).

5. Eiraku, M. et al. Self-organizing optic-cup morphogenesis in three-dimensional culture. Nature 472, 51–56 (2011).

6. Meinhardt, A. et al. 3D Reconstitution of the Patterned Neural Tube from Embryonic Stem Cells. Stem Cell Rep. 3, 987–999 (2014).

7. Martyn, I., Kanno, T. Y., Ruzo, A., Siggia, E. D. & Brivanlou, A. H. Self-organization of a human organizer by combined Wnt and Nodal signaling. Nature 558, 132–135 (2018).

8. Bikoff, J. B. et al. Spinal Inhibitory Interneuron Diversity Delineates Variant Motor Microcircuits. Cell 165, 207–219 (2016).

9. Sagner, A. & Briscoe, J. Establishing neuronal diversity in the spinal cord: A time and a place. Development (Cambridge) vol. 146 (2019).

10. Dessaud, E., McMahon, A. P. & Briscoe, J. Pattern formation in the vertebrate neural tube: a sonic hedgehog morphogen-regulated transcriptional network. Development 135, 2489–2503 (2008).

11. Le Dréau, G. & Martí, E. Dorsal-ventral patterning of the neural tube: A tale of three signals. Dev. Neurobiol. 72, 1471–1481 (2012).

12. Jessell, T. M. Neuronal specification in the spinal cord: inductive signals and transcriptional codes. Nat. Rev. Genet. 1, 20–29 (2000).

13. Yamada, T., Placzek, M., Tanaka, H., Dodd, J. & Jessell, T. M. Control of cell pattern in the developing nervous system: Polarizing activity of the floor plate and notochord. Cell 64, 635–647 (1991).

14. Patten, I., Kulesa, P., Shen, M. M., Fraser, S. & Placzek, M. Distinct modes of floor plate induction in the chick embryo. Development 130, 4809–4821 (2003).

15. Placzek, M., Tessier-Lavigne, M., Yamada, T., Jessell, T. & Dodd, J. Mesodermal Control of Neural Cell Identity: Floor Plate Induction by the Notochord. Science 250, 985–988 (1990).

16. Placzek, M. & Briscoe, J. The floor plate: multiple cells, multiple signals. Nat. Rev. Neurosci. 6, 230–240 (2005).

17. Kremnyov, S., Henningfeld, K., Viebahn, C. & Tsikolia, N. Divergent axial morphogenesis and early shh expression in vertebrate prospective floor plate. EvoDevo 9, 1–17 (2018).

18. Ranga, A. et al. Neural tube morphogenesis in synthetic 3D microenvironments. Proc. Natl. Acad. Sci. 113, (2016).

19. Ishihara, K. et al. Topological morphogenesis of neuroepithelial organoids. Nat. Phys. (2022) doi:10.1038/s41567-022-01822-6.

20. Sasaki, H., Hui, C., Nakafuku, M. & Kondoh, H. A binding site for Gli proteins is essential for *HNF-3* β floor plate enhancer activity in transgenics and can respond to Shh in vitro. Development 124, 1313–1322 (1997).

21. Etoc, F. et al. A Balance between Secreted Inhibitors and Edge Sensing Controls Gastruloid Self-Organization. Dev. Cell 39, (2016).

22. Stafford, D. A., Brunet, L. J., Khokha, M. K., Economides, A. N. & Harland, R. M. Cooperative activity of noggin and gremlin 1 in axial skeleton development. Development 138, 1005–1014 (2011).

23. Arnold, K. et al. Sox2+ Adult Stem and Progenitor Cells Are Important for Tissue Regeneration and Survival of Mice. Cell Stem Cell 9, 317–329 (2011).

24. Balmer, S., Nowotschin, S. & Hadjantonakis, A.-K. Notochord morphogenesis in mice: Current understanding & open questions: Notochord Morphogenesis in Mice. Dev. Dyn. 245, 547–557 (2016).

25. Turing, A. M. The chemical basis of morphogenesis. Philos. Trans. R. Soc. Lond. B. Biol. Sci. 237, 37–72 (1952).

26. Gierer, A. & Meinhardt, H. A theory of biological pattern formation. Kybernetik 12, 30– 39 (1972).

27. Zheng, Y. et al. Dorsal-ventral patterned neural cyst from human pluripotent stem cells in a neurogenic niche. Sci. Adv. 5, eaax5933 (2019).

28. Abdel Fattah, A. R., et al. Actuation enhances patterning in human neural tube organoids. Nat. Commun. 12, (2021).

29. Abdel Fattah, A. R., et al. Neuroepithelial Organoid Patterning is Mediated by Wnt-Driven Turing Mechanism. bioRxiv (2021) doi:10.2139/ssrn.3811873.

30. Lancaster, M. A. et al. Cerebral organoids model human brain development and microcephaly. Nature 501, 373–379 (2013).

31. Van Den Brink, S. C., et al. Symmetry breaking, germ layer specification and axial organisation in aggregates of mouse embryonic stem cells. Development 141, 4231–4242 (2014).

32. Takata, N., Sakakura, E., Eiraku, M., Kasukawa, T. & Sasai, Y. Self-patterning of rostral-caudal neuroectoderm requires dual role of Fgf signaling for localized Wnt antagonism. Nat. Commun. 8, 1339 (2017).

33. Rivron, N. C. et al. Blastocyst-like structures generated solely from stem cells. Nature 557, (2018).

34. Turner, D. A., Baillie-Johnson, P. & Martinez Arias, A. Organoids and the genetically encoded self-assembly of embryonic stem cells. BioEssays 38, 181–191 (2016).

35. McNamara, H. M., Solley, S. C., Adamson, B., Chan, M. M. & Toettcher, J. E. Recording morphogen signals reveals origins of gastruloid symmetry breaking. http://biorxiv.org/lookup/doi/10.1101/2023.06.02.543474 (2023) doi:10.1101/2023.06.02.543474.

36. Suppinger, S. et al. Multimodal characterization of murine gastruloid development. Cell Stem Cell 30, 867–884.e11 (2023).

37. Simunovic, M. et al. A 3D model of a human epiblast reveals BMP4-driven symmetry breaking. Nat. Cell Biol. 21, 900–910 (2019).

38. Shao, Y. et al. A pluripotent stem cell-based model for post-implantation human amniotic sac development. Nat. Commun. 8, 208 (2017).

39. Mori, S. et al. Self-organized formation of developing appendages from murine pluripotent stem cells. Nat. Commun. 10, 3802 (2019).

40. Serra, D. et al. Self-organization and symmetry breaking in intestinal organoid development. Nature 569, 66–72 (2019).

41. Le Dréau, G. et al. Canonical BMP7 activity is required for the generation of discrete neuronal populations in the dorsal spinal cord. Development 139, 259–268 (2012).

42. Lee, K. J., Dietrich, P. & Jessell, T. M. Genetic ablation reveals that the roof plate is essential for dorsal interneuron specification. Nature 403, 734–740 (2000).

43. Lee, K. J., Mendelsohn, M. & Jessell, T. M. Neuronal patterning by BMPs: a requirement for GDF7 in the generation of a discrete class of commissural interneurons in the mouse spinal cord. Genes Dev. 12, 3394–3407 (1998).

44. Dale, K. et al. Differential patterning of ventral midline cells by axial mesoderm is regulated by BMP7 and chordin. Development 126, 397–408 (1999).

45. McMahon, J. A. et al. Noggin-mediated antagonism of BMP signaling is required for growth and patterning of the neural tube and somite. Genes Dev. 12, 1438–1452 (1998).

46. Furuta, Y., Piston, D. W. & Hogan, B. L. M. Bone morphogenetic proteins (BMPs) as regulators of dorsal forebrain development. Development 124, 2203–2212 (1997).

47. Burtscher, I., Barkey, W. & Lickert, H. Foxa2-venus fusion reporter mouse line allows live-cell analysis of endoderm-derived organ formation: Foxa2-Venus Fusion Reporter. genesis 51, 596–604 (2013).

48. Chu, V. T. et al. Efficient generation of Rosa26 knock-in mice using CRISPR/Cas9 in C57BL/6 zygotes. BMC Biotechnol. 16, 4 (2016).

49. Ran, F. A. et al. Genome engineering using the CRISPR-Cas9 system. Nat. Protoc. 8, 2281–2308 (2013).

50. Matsumoto, K. et al. Advanced CUBIC tissue clearing for whole-organ cell profiling. Nat. Protoc. 14, 3506–3537 (2019).

51. Haase, R. et al. CLIJ: GPU-accelerated image processing for everyone. Nat. Methods 17, 5–6 (2020).

52. Stirling, D. R. et al. CellProfiler 4: improvements in speed, utility and usability. BMC Bioinformatics 22, 433 (2021).

53. Berg, S. et al. ilastik: interactive machine learning for (bio)image analysis. Nat. Methods 16, 1226–1232 (2019).

54. Mitchell, N. P. & Cislo, D. J. TubULAR: Tracking in toto deformations of dynamic tissues via constrained maps. http://biorxiv.org/lookup/doi/10.1101/2022.04.19.488840 (2022) doi:10.1101/2022.04.19.488840.

55. Rodolà, E., Bulò, S. R. & Cremers, D. Robust Region Detection via Consensus Segmentation of Deformable Shapes. Comput. Graph. Forum 33, 97–106 (2014).

56. Crane, K., Livesu, M., Puppo, E. & Qin, Y. A Survey of Algorithms for Geodesic Paths and Distances. (2020) doi:10.48550/ARXIV.2007.10430.

57. Pallares, L. F., Picard, S. & Ayroles, J. F. TM3’seq: A Tagmentation-Mediated 3’ Sequencing Approach for Improving Scalability of RNAseq Experiments. G3 GenesGenomesGenetics 10, 143–150 (2020).

58. Truett, G. E. et al. Preparation of PCR-Quality Mouse Genomic DNA with Hot Sodium Hydroxide and Tris (HotSHOT). BioTechniques 29, 52–54 (2000).

